# A spatio-temporal brain miRNA expression atlas identifies sex-independent age-related microglial driven miR-155-5p increase

**DOI:** 10.1101/2025.03.15.643430

**Authors:** Annika Engel, Viktoria Wagner, Oliver Hahn, Aulden G. Foltz, Micaiah Atkins, Amila Beganovic, Ian H. Guldner, Nannan Lu, Aryaman Saksena, Ulrike Fischer, Nicole Ludwig, Eckart Meese, Tony Wyss-Coray, Andreas Keller

## Abstract

An in-depth understanding of the molecular processes composing aging is crucial to develop therapeutic approaches that decrease aging as a key risk factor for cognitive decline. Herein, we present a spatio-temporal brain atlas (15 different regions) of microRNA (miRNA) expression across the mouse lifespan (7 time points) and two aging interventions composed of 1009 samples. MiRNAs are promising therapeutic targets, as they silence genes by complementary base-pair binding of messenger RNAs and are known to mediate aging speed. We first established sex- and brain-region-specific miRNA expression patterns in young adult samples. Then we focused on sex-dependent and independent brain-region-specific miRNA expression changes during aging. The corpus callosum in males and the choroid plexus in females exhibited strong sex-specific age-related signatures. In this work, we identified three sex-independent brain aging miRNAs (miR-146a-5p, miR-155-5p and miR-5100). We showed for miR-155-5p that these expression changes are driven by aging microglia. MiR-155-5p targets mTOR signaling pathway components and other cellular communication pathways and is hence a promising therapeutic target.

## Introduction

Aging which is defined as the loss of physiological functions resulting in the end of lifespan impacts all organs^1^. But especially the mechanisms driving the gradual decline in the brain and the resulting loss of its functionalities are yet to be understood^2^. Previous molecular and neuroimaging studies suggested that the different regions of the brain may experience varying degrees of susceptibility to the effects of the aging process^3,4^. Brain aging atlases of epigenetic modifications^5^ and transcriptional changes^3^ were previously generated, but the miRNA expression layer is still missing. Mature miRNAs are short (18-24 nt) single stranded RNA molecules which play an important role in post-transcriptional regulation via a complementary base pair binding to mRNA molecules^6^, thereby reducing their stability and inhibiting translation^7^. Furthermore, their capacities of targeting multiple mRNAs in the same pathways^8^ as well as their stability make them promising targets for therapy applications and diagnostics^9^. MiRNAs have been identified as mediators of protective microglial states in an AD mouse model^10^, proposed as important drivers of aging-related phenotypes cross-organs^11^ and been shown to control aging speed via exosomes^12^. Hence, generating a comprehensive atlas of miRNA expression during healthy brain aging is urgently needed to characterize how and where aging occurs and leads to vulnerabilities, such as the development of neurological disorders like Alzheimer’s or Parkinson’s disease^13–17^. Adding to existing mRNA and protein datasets, miRNA expression data will increase our understanding of the complex multilayered processes that lead to the heterogeneous aging phenotype. Currently the information on expression patterns of miRNAs in different brain regions is rather sparse^18–20^. Many studies focus on a set of 2-12 regions and predominantly used male mice with few to no differing age groups^18,20^. Furthermore, through NGS techniques we were able to assess another layer of miRNA biology, previously unexplored in the aging context, the isomiR expression. IsomiRs are naturally occurring variants of the mature archetype miRNA and are clinically relevant as well^21^.

We decided to use inbred mouse samples from C57BL/6JN and thereby minimize genetic as well as environmental variability to create a baseline of miRNA expression changes with minimal confounding factors. A baseline derived from a model organism is essential for human studies as these are challenging due to sparse availability of unaffected brains, genetic and environmental heterogeneity, varying post-mortem intervals and hence varying sample quality. In our study, we do not only aim to understand temporally resolved expression changes in different regions by establishing a wide and sex-specific atlas of brain-region-specific miRNA expression patterns. We analyze region-specific miRNA expression patterns in males and females separately to uncover sex-specific regional expression. Building on the sex-specific analysis, we study age-related miRNA expression changes in a sex dependent and independent manner to identify common and specific signatures. We found three cross-sex brain aging miRNAs, likely derived from microglial expression changes. Hence, we collected young and aged microglia to analyze their miRNA expression. We checked the expression changes of our brain aging miRNAs in two commonly used aging interventions, namely, dietary restriction and young plasma injection. Finally, we combined our miRNA data set with the existing mRNA dataset and found cellular communication and mTOR signaling pathways regulated by the cross-sex brain aging miRNA miR-155-5p.

## Results

### Brain region-specific miRNA expression patterns

Aging is the main risk factor to suffer from major neurodegenerative diseases, such as Alzheimer’s disease (AD) and Parkinson’s disease (PD) as well as cognitive dysfunction. Therefore, understanding the underlying mechanisms is crucial for the development of effective therapies^13^. We generated bulk sequencing data from 844 samples from 15 defined regions at seven age stages (3, 12, 15, 18, 21, 26, 28 months) to uncover the region-specific miRNA expression patterns during aging. Samples from the following regions were collected: corpus callosum, choroid plexus, neurogenic subventricular zone (SVZ), hippocampus anterior and posterior, hypothalamus, thalamus, caudate putamen, pons, medulla, cerebellum, olfactory bulb and three cortical regions, namely, motor, entorhinal and visual cortex (Fig. 1a). To allow for the highest data quality, we ensured that all samples included in the analysis carry at last 2 million reads aligned to the mouse genome. On average we found a read count of over 9.5 million per group, when aggregating all samples according to combinations of brain region and age (Supplementary Fig. 1a). This approach ensured that, on average, over 55% of the reads were mapped to miRNAs (Supplementary Fig. 1b). In total 828 of 844 samples (98%) passed our strict quality control (Supplementary Table 1 and 2). In addition to sample filtering, we conducted a feature filtering ensuring that at minimum 10% of the samples within at least one brain region exceeded a raw count of 5. If a RNA fulfilled this requirement, we considered it expressed within that brain region. A subsequent UMAP visualization of the features (miRNAs, lncRNAs, piRNAs, rRNAs, scaRNAs, snoRNAs, snRNAs and tRNAs) showed a distinct separation of several brain regions, among those the olfactory bulb and the cerebellum were the most distinctly different ones (Supplementary Fig. 1c). Neither sex nor age can be identified as strong factors driving a grouping (Supplementary Fig. 1d and 1e). Analyzing the composition of expressed counts per brain region indicated a homogenous distribution (Supplementary Fig. 1f). Within most brain regions among these cerebellum, motor cortex, hippocampus anterior and olfactory bulb, the composition of expressed counts for all RNA classes remains at a mainly constant level over all ages (Supplementary Fig. 1g). Correlating each feature with age per brain region revealed 720 positive correlated features for tRNAs and 127 negatively correlated ones for miRNAs (51 positively correlated miRNAs, Fig. 1b). We exemplary highlight a tRNA (tRNA-Glu-TTC-1-1), which was significantly positively correlated with age in both male and female in multiple brain regions (Fig. 1c). Motivated by these high feature numbers exhibiting a correlation with age, we visualized UMAP results of 404 tRNAs which failed to provide a distinct separation into brain regions, sex or age (Fig. 1d, Supplementary Fig. 2a and 2b). This circumstance is strengthened by a PVCA which explains 36% of the observed variance with the brain region identity (Fig. 1e). The miRNAs in contrast, exhibit a larger share of 54% of the explained variance by brain region which reflects in the clear separation in the UMAP visualization (Fig. 1f and 1g). Yet, neither age nor sex can be identified as a driving factor for miRNAs (Supplementary Fig. 2c and 2d). Age, as an independent factor, introduced 0.6% of the overall variation and 3.8% in combination with the brain region andthe variable sex was responsible for 0.2% of the overall variation and for 2.8% in combination with the brain region (Fig. 1g). In total, we included 1,174 miRNAs, of which 785 were expressed across all brain regions (Supplementary Fig. 2e). Since our sequencing protocol is optimized for miRNAs and we found the greatest region-specificity within miRNAs and a high number of age-correlated ones we focused the following analysis exclusively on miRNAs. An overview of our miRNA expression data can be found at https://ccb-compute2.cs.uni-saarland.de/brainmirmap.

**Fig. 1:**
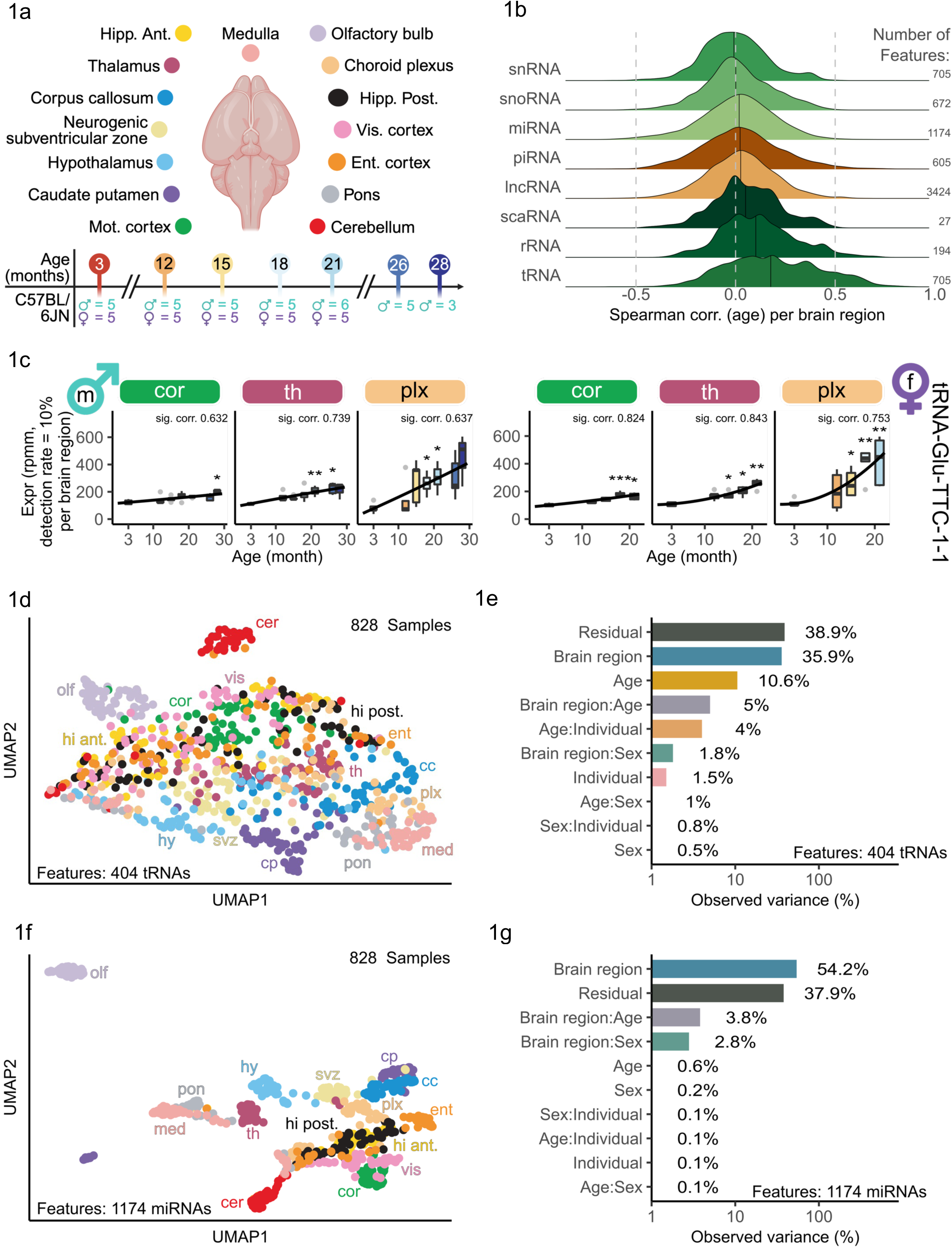
Spatio-temporal overview and analysis of RNA expression in the aging mouse brain. **a** Study overview: Age stages of tissue collection at 3, 12, 15, 18, 21, 26 and 28 months in 15 different brain regions defined according to the Allen Brain atlas. The numbers below the timeline are indicating how many mouse individuals were observed per age stage and sex. Schematic plot created with BioRender.com. **b** The distribution of Spearman’s rank correlation coefficients between age and individual features for each RNA class. These correlations were calculated separately for each brain region using all available samples. The black solid line within each ridgeline indicates the median. On the right we see the number of features included in the ridgeline. **c** Boxplots showing the time series of tRNA tRNA-Glu-TTC-1-1 for three different brain regions each for male left and female right. The Spearman’s rank correlation coefficient from the miRNA expression with age is displayed above each plot. An asterisk indicates the significance level. **d** UMAP of all samples from all regions and for all tRNA features colored by regions as indicated in Fig. 1a. **e** Principal Variance Component Analysis for the tRNAs over all samples for the biological factors brain region, age, sex, and individual depicting the observed variance in percent. **f** UMAP of all samples from all regions and for all miRNA features colored by regions as indicated in Fig. 1a. **g** Principal Variance Component Analysis for the miRNAs again over all samples for the biological factors brain region, age, sex, and individual depicting the observed variance in percent. Colours are matching to the once from Fig. 1e.

We herein present, to our knowledge, the first study analyzing miRNA expression in a spatial resolution of 15 regions within the male and female brain using high-throughput sequencing. Previously only micro-array-based studies were performed with up to 13 regions in male mice only^18,19^. Using NGS methods in addition to having a greater sample size of each sex enabled us to identify more expressed miRNAs compared to microarray-based techniques. Furthermore, with high-throughput sequencing isomiR alterations of miRNAs can be identified, which is not possible in microarray-based studies.

First, we hence focused on identifying strong region-specific signatures of miRNA expression in the different brain regions for both sexes unified and individually. Therefore, we analyzed the expression of young adult mice (ages: 3, 12 and 15 months) without considering the age effect. Visualizing the UMAP result for the young adult mice exhibited a clear separation driven by the brain region identity (Supplementary Fig. 2f). Exceptions like the overlap of hippocampus anterior and posterior can be explained by their anatomical and functional proximity. The closeness of SVZ, corpus callosum and caudate putamen reflects their anatomical proximity and show consistent sampling precision over all mice. Neither sex nor age can be identified as strong factors driving a distinct grouping in the UMAP visualization in young only samples (Supplementary Fig. 2g and 2h).

To identify miRNAs driving the region-specific grouping (Supplementary Fig. 2f), we selected 50 miRNAs according to the highest coefficient of variation (Supplementary Fig. 2i). We identified miRNAs which were distinctly expressed with respect to the brain average (highlighted with a black border) and clustered the respective brain regions into four clusters (cf. Methods). The strongest region-specific signature was observed in the olfactory bulb, since it formed a separate cluster. Fifteen miRNAs were distinctly expressed with respect to the brain average. Among these miRNAs were miR-200a-5p, miR-200a-3p, miR-200b-5p, miR-200b-3p and miR-200c-5p. MiR-200 family members have been reported in the olfactory bulb as crucial to mediate neuronal maturation through targeting *Zeb2* during postnatal development^22^.

Splitting the data into male and female samples however revealed differing signatures compared to the ones observed in the combined data (Fig. 2a), underlining the importance of a sex-separable data set. For the male samples, the strong unique signature of the olfactory bulb persisted, whereas for the females this distinct signature is less pronounced. We still observed distinct expression of members of the miR-200 family (miR-200b/c-5p). In females, pons and medulla also exhibited distinct regional expression signatures, by each forming a cluster by themselves. Especially interesting were the distinctly expressed signature miRNAs in the medulla, miR-10b-3p and miR-10b-5p, and shared between medulla and pons miR-1a-1-5p and miR-10a-3p/5p. Since miR-1 and miR-10b were previously reported to regulate BDNF (brain-derived neurotrophic factor) an important protein involved in synaptogenesis, memory and leaning^23^. In males, a strict anatomical organization of miRNA expression was observed as the olfactory bulb anterior had a distinct profile differing from the central brain regions (corpus callosum, subventricular zone, hypothalamus, thalamus, choroid plexus) and the posterior regions medulla, pons and cerebellum were dominated by another set of miRNAs.

**Fig. 2:**
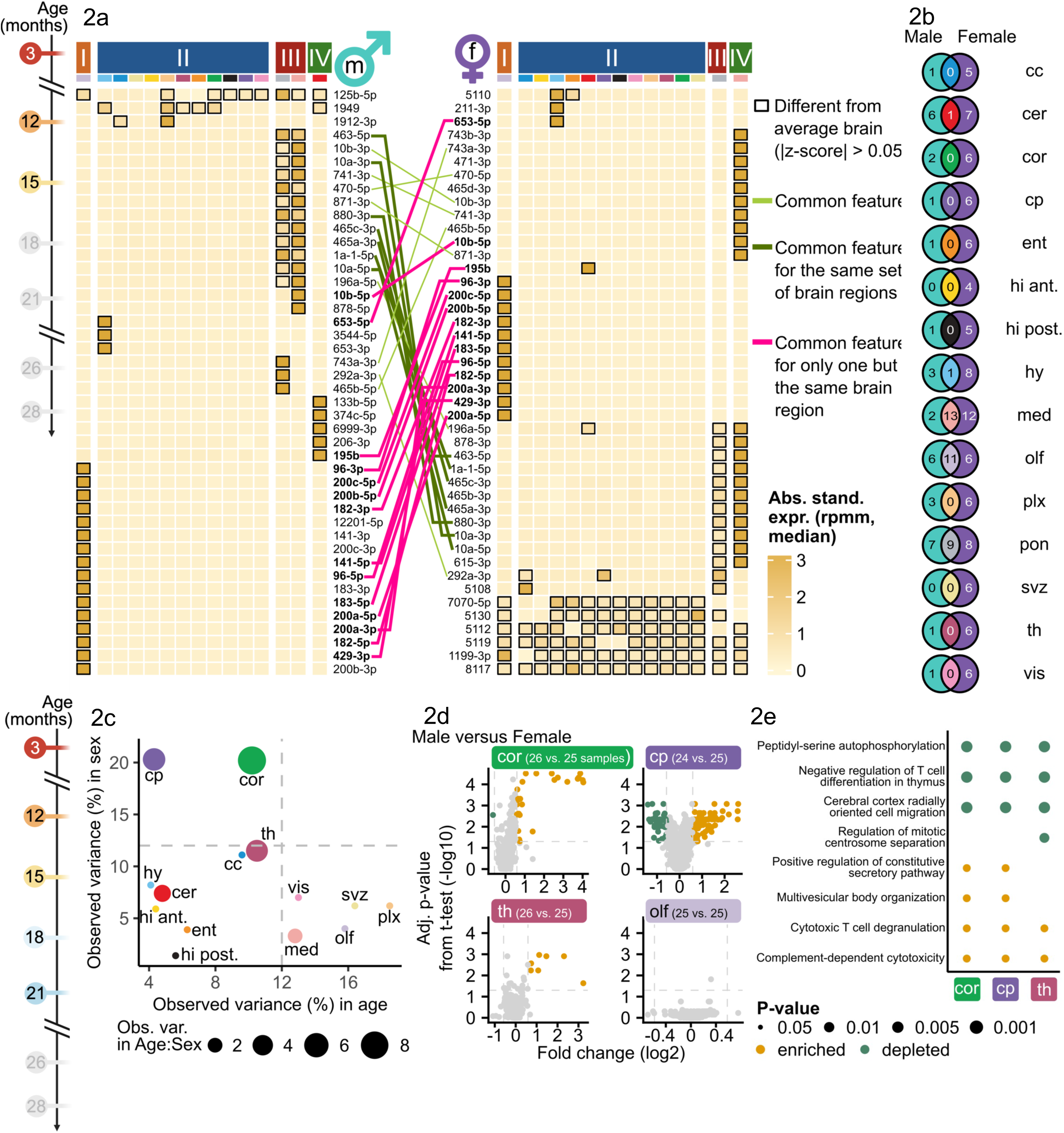
Sex- and brain region-specific miRNA variation in the mouse brain. **a** Heatmaps of the 50 top miRNA from all brain regions determined by coefficient of variation calculated using the medians of the expression values of each brain region. Shown are the absolute standardized expression values (z-scores) for the younger male (left) and younger female (right) samples. The black borders are indicating the binarization (|z-score| > 0.5) on which a clustering into four clusters using a hierarchical clustering was applied. We consider a miRNA in a brain region different from the average brain if it surpasses the before mentioned threshold. If it only exceeds the threshold for one brain region, we call it brain region specific. The lines connecting the two heatmaps are highlighting the occurrence of common features (light green). If features are differing from the average brain for only one but the same brain region in both sexes, they are highlighted in pink and in dark green if they differ from the average brain in the same set of multiple brain regions. For visualization purposes, we removed features entirely below the selected threshold. **b** Venn diagram linking the features which are differing from the average brain per brain region between the male and the female heatmap. **c** Principal Variance Component Analysis showing the observed variance for each brain region individually over all sex-matched samples (3, 12, 15, 18, 21 months) for the biological factors age and sex. The point size indicates the variance when observing age and sex in combination. Combined variance shares are indicated in point size. The colors refer to the regions from A. Thresholds at 12% for both axes are marked with grey dashed lines. **d** Volcano plot for the sex-specific comparison of the brain regions mot. cortex, choroid plexus, thalamus and olfactory bulb. Colored dots indicate significantly down (green) and significantly upregulated (yellow) miRNAs (fold change ≥ 1.5 or ≤ 1/1.5, adjusted p-value < 0.05). **e** The gene set enrichment analysis (GSEA) result obtained from MIEAA^54^ for three brain regions showing the top 10 (sorted adjusted by p-value) depleted (green) and enriched (yellow) pathways over the three brain regions.

Summarizing these results, we determined sex-unspecific miRNA regional expression patterns (Fig. 2b, Supplementary Table 3). In medulla, we observed thirteen sex-unspecific miRNAs (e.g. miR-10a/b), eleven in the olfactory bulb (e.g. miR-200b/c-5p), nine in pons, miR-195b in cerebellum and miR-653-5p in hypothalamus. In contrast, we identified sex-specific miRNAs with regional expression patterns in thirteen regions for males (e.g. miR-133b-5p in cerebellum) and in all regions for females (e.g. miR-471-3p in medulla). Hence, we concluded that regional miRNA expression patterns occur in a sex independent manner in anterior and posterior brain regions (olfactory bulb, cerebellum, medulla, pons) and strong sex-specific regional patterns rather occur in central brain regions (corpus callosum, motor cortex, choroid plexus, thalamus).

### Sex as a factor for miRNA expression during aging expression patterns

Considering the importance of sex as a variable in regional expression patterns, we chose to investigate the impact of sex as a variable within our dataset compared to age. Within this dataset, we collected both male and female samples until the age of 21 months. During sex-specific analysis, we were limited to 14 brain regions due to an insufficient number of samples from older female mice in pons.

Region-wise analysis of variation shares of age, sex and their combination revealed that for two regions -mot. cortex and caudate putamen -the observed variation introduced by the variable sex was 20.2% and 20.3%, respectively (Fig. 2c). Whereas the variation share of age and their combination was under 12% for both regions. In particular for the caudate putamen, age accounted only for 4.3% of the observed variation. Sex differences in pathologies as well as neural properties and associated mechanisms have been reported previously for this region^24^, e.g. in humans for fiber connection strength.

We identified 86 significantly upregulated and 37 significantly downregulated miRNAs between male and female in the caudate putamen (fold change greater or equal than 1.5 or smaller or equal than 1/1.5, adjusted p-value smaller than 0.05) (Fig. 2d). Amongst the top three significantly upregulated miRNAs according to fold change were miR-128-1-5p, miR-490-3p and miR-344b-3p. In the motor cortex, 26 miRNAs were significantly upregulated and one miRNA significantly downregulated. Thalamus and corpus callosum also exhibited solely sex-driven variances above 10% but equally strong solely age-driven variances. In the thalamus, however, only seven miRNAs were significantly upregulated between male and female. In contrast, regions with a low share of variation explained by sex, such as olfactory bulb exhibited no significantly differentially expressed miRNAs between males and females. A gene set enrichment analysis over the miRNAs ranked according to their significance and sex-deregulation in mot. cortex, caudate putamen and thalamus revealed overlaps between the regulated pathways (Fig. 2e). E.g. “cerebral cortex radially oriented cell migration” was depleted in all three regions and “complement-dependent cytoxicity” was enriched. In summary, we found sex-specific miRNA expression patterns that persist during the entire lifespan and dominate over age-related expression changes, especially strong in motor cortex and caudate putamen. These changes potentially relate to sex-specific regulation of distinct pathways.

However, our study focusses on age-related miRNA expression patterns and for five regions the share of variation explainable by age (> 12%) dominated over the share explainable by sex (< 10%) and the combination between age and sex (< 3%) (Fig. 2c). These regions, namely, subventricular zone, olfactory bulb, vis. cortex and medulla and most prominently choroid plexus (18.5% of variation explainable by age) were especially interesting for the main research question whether age-related miRNA expression changes in the brain occur region specifically.

### Sex-specific miRNA expression changes during aging

As we observed differing expression patterns between male and female in our initial analysis, we first chose to analyze male and female samples separately to verify common and sex-specific effects. By collecting multiple discrete age stages in our data set, we were able to approximate the aging trajectories over the lifespan for each miRNA in each region and for both sexes. Analyzing trajectories is crucial as expression changes can occur in differing degrees and varying shapes, but in similar directions. These details remain hidden in binary comparisons of young versus aged samples. Therefore, our data set consists of seven age stages spread over the lifespan of a mouse. However, considering age as a discrete variable may still underestimate the impact of aging, since it is continuous by nature.

As miR-9 family members were previously extensively studied in the brain and showed brain-specific expression^25–27^, we exemplarily investigate miR-9 expression in our data set. Considering all samples miR-9-5p is highly expressed in all regions (Supplementary Fig. 2j) though it is not amongst the 50 most variable miRNAs for the young samples according to the coefficient of variation (Supplementary Fig. 2i). In the olfactory bulb, we detected a median expression of over 100k rpmm. In contrast, second highest expression in motor cortex was around 48k and in medulla the median expression was below 20k. During aging the miR-9-5p expression remained mostly stable in medulla and motor cortex in both sexes (Fig. 3a, Supplementary Fig. 3a). However, the expression in the olfactory bulb slightly decreased over the lifespan in males but remained mostly stable in females. MiR-9-5p is known to be involved in neurogenesis, axon development, differentiation and proliferation of neural progenitor cells^27^. Hence, further investigation of functional consequences of its region-specific and sex-specific aging expression could yield to additional insights into these mechanisms.

**Fig. 3.**
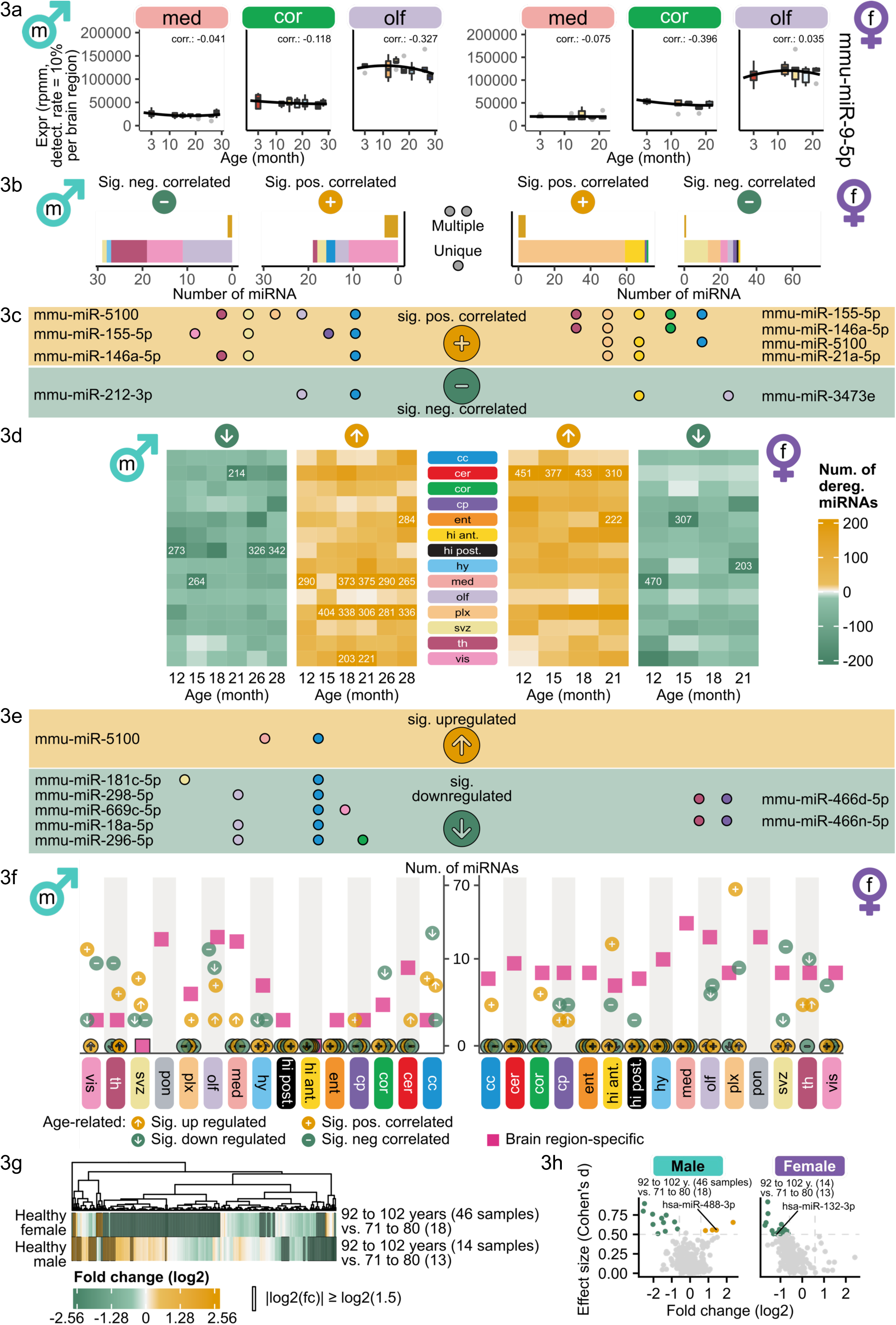
Sex-specific miRNA dynamics in brain aging across male and female. Sex-specific analysis. We split the figure in a left and a right part covering the male and female samples, respectively. **a** Boxplots showing the time series of miRNA mmu-miR-9-5p for three different brain regions each. The Spearman’s rank correlation coefficient from the miRNA expression with age is displayed above each plot. **b** Bar plots showing the number of significantly anti- or positively correlated miRNAs with age per brain region (using Spearman’s rank correlation coefficient with |R| ≥ 0.5, adjusted p-value < 0.05). The upper bars contain miRNAs which are in the same direction significantly correlated in more than one brain region (in yellow). The respective lower bars contain miRNAs that are unique for one brain region (colored in the corresponding brain region color). **c** List of features which are significantly correlated for more than one brain region. The dots indicate the brain region(s) (colored according to Fig. 1a) in which the feature is significantly positively correlated (yellow; top) and significantly anti-correlated (green; bottom). **d** Heatmaps showing for each brain region the number of up- or deregulated miRNAs for the comparison between each age stage to 3 months (fold change ≥ 1.5 or ≤ 1/1.5, adjusted p-value < 0.05). Numbers in cell displayed when above threshold of 200 miRNAs. **e** List of features which are in the same direction significantly deregulated for at least one age comparison within at least two brain regions with significantly upregulated miRNAs at the top and significantly downregulated ones at the bottom. Analogous to Fig. 3c, the colors of the dots indicate the brain region in which the deregulations can be found. **f** Shows the amount of brain region-specific and age-related features per approach. Purple markers indicate the number of brain region-specific features from Fig. 2a, arrows pointing to the top (bottom) show the amount of significantly upregulated (downregulated) miRNAs in at least one age comparison and positive (negative) signs show the amount of significantly positively (anti-) correlated miRNAs. Human miRNA data from ROSMAP^29^: **g** We divide the healthy samples into three similar sized sets according to their age of death (71 to 80, 81 to 91 and 92 to 102 years). Using the youngest and the oldest group we perform a DE analysis per sex. The color denotes the fold change for the feature (columns) and the black border if the fold change is greater or equal than 1.5 or smaller or equal than 1/1.5. A star would indicate a significance based on the adjusted p-value (Student’s *t*-test, Benjamini-Hochberg procedure). **h** Scatter plot of the Cohen’s d values against the log2-normalized fold changes colored if exceeding the thresholds for the fold change (|log2(fc)| ≥ log2(1.5)) and of the effect size (|d| ≥ 0.5). Yellow for an upregulation and green in case of a downregulation. The labeled miRNAs are those which were already identified as age-correlated for the brain region mot. cortex (Supplementary Fig. 4c).

We measured the relation of each miRNA with age in each brain region by calculating the Spearman’s rank correlation coefficient in males and females to identify sex-region-specific aging miRNAs. MiRNAs with correlation values exceeding the interval between −0.5 and 0.5 were deemed negatively (anti-correlated) or positively correlated with age, respectively. Across all brain regions, we observed significantly correlations with age for 13.79% of the miRNAs in males and 37.87% in females (Supplementary Fig. 3b). In males, twelve miRNAs were significantly positively correlated with age in the vis. cortex and in olfactory bulb twelve miRNAs were significantly anti-correlated (Fig. 3b, Supplementary Table 4). In females, 63 miRNAs were significantly positively correlated in choroid plexus and 13 were significantly anti-correlated in the SVZ. Three miRNAs in males and four miRNAs in females were significantly positively correlated with age in more than three brain regions (Fig. 3c). MiR-212-3p was significantly anti-correlated with age in males and miR-3473e in females both for two brain regions namely olfactory bulb and corpus callosum, hippocampus anterior and olfactory bulb, respectively. MiR-155-5p, miR-146a-5p and miR-5100 were significantly positively correlated with age in males and females, exhibiting a strong aging signature independent from sex and region. Additionally, miR-21a-5p was significantly positively correlated with age in hippocampus anterior and choroid plexus in females. Given that both miR-155-5p and miR-146a-5p are established regulators of neuroinflammation and are implicated in neurodegenerative diseases^28^, their roles in brain aging present compelling targets for further investigations.

By calculating correlations, we neglected non-monotonic effects in the trajectories. Therefore, we additionally considered the differences between the older ages (12 to 28 months for male and 12 to 21 months for female) and the first age stage (3 months) by performing a differential expression (DE) analysis using these comparisons (Supplementary Table 5). We count a miRNA as upregulated (downregulated) in a brain region if at least one age comparison within the brain region exhibits a fold change greater or equal than 1.5 (smaller or equal than 1/1.5).

In the choroid plexus in males, exclusively one miRNA (mmu-miR-5100) was significantly positively correlated with age. However more than 280 miRNAs were upregulated at 15 months and all later time points in males in this region (Fig. 3d). Out of these 280 miRNAs 204 miRNAs were consistently upregulated in all consecutive time points from 15 to 28 months. These results indicate that the expression of these miRNAs is drastically increased between 12 and 15 months and remained constantly high thereafter, which was not identified during correlation analysis. In females, we overall observed 76 significantly positively age-correlated miRNAs (Fig. 3b) and a matching high count of upregulated miRNAs is observed within the age comparisons across all regions (Fig. 3d). Especially the brain regions cerebellum and choroid plexus stood out with 511 and 308 distinct miRNAs over all age comparisons, respectively. While cerebellum exhibited an even higher upregulation trend in females starting at 12 months of age with 451 upregulated miRNAs, no significant age-correlated miRNAs were observed before.

Though we observed strong deregulation trends for many miRNAs in different regions, few were significantly deregulated (Supplementary Fig. 3c). Therefore, we aggregated the miRNAs that were significantly deregulated uniquely and in multiple regions to determine the strongest age-deregulated miRNAs per sex (Supplementary Fig. 4a). MiR-5100, also identified as a significant age-correlated miRNA in multiple brain regions in both sexes (Fig. 3c), was significantly upregulated in males in the medulla and corpus callosum (Fig. 3e), with no significant regulation observed in females in any region. More miRNAs were significantly downregulated in unique regions in males than in females (Supplementary Fig. 4a).

We aggregated the miRNAs obtained from the age-correlation and from the deregulation analysis to a candidate set presenting all the results from the age-related analysis regardless their direction (Supplementary Fig. 4b, Supplementary Table 6). In males, corpus callosum, visual cortex and olfactory bulb were the most prominent regions affected by miRNA expression changes as opposed to choroid plexus in females. Connecting all previous analysis, we summarized the region-specific and age-related signatures in each sex (Fig. 3f). In males, the olfactory bulb showed the highest number of region-specific miRNAs. Even though there is also a high number of significant age-related miRNAs, we found no region-specific miRNA that also exhibited strong age-related expression changes in any region. In females, we detected region-specific miRNAs as well as significant age-related ones but again no simultaneously region-specific and age-related miRNAs were detected. Our findings indicate that miRNAs exhibiting a strong regionally defined expression pattern in the brain have a stable expression during aging. In contrast, even though age-related miRNAs occur in a region dependent manner, these miRNAs were not amongst the highest variable ones between all regions.

To see whether miRNA candidates from the sex-specific bulk mouse data showed similar trends in humans, we investigated miRNA expression patterns within the ROSMAP dataset^29^. This dataset includes 203 healthy patients (71 males and 132 females). We split the ages at time of death into three groups (71 to 80, 81 to 91 and 92 to 102 years). Between the oldest and the youngest group, we conducted a DE analysis. In females, we obtained 144 downregulated (fold change smaller or equal than 1/1.5) and ten upregulated miRNAs (greater or equal than 1.5) and 42 downregulated and 32 upregulated miRNAs in males (Fig. 3g). None of these are significant according to the adjusted Student’s *t*-test p-values using Benjamini-Hochberg procedure for adjustment and a significance level of 0.05. Elven miRNAs were downregulated and four were upregulated in males and obtained an abolute effect size greater or equal than 0.5 calculated using Cohen’s d (Fig. 3h). Whereas in females 19 downregulated miRNAs exceeded the effect size threshold and no upregulated ones. Considering these miRNAs as age-related candidates in the human dataset, we intersected these with miRNAs found as age-correlated in mot. cortex, the closest matching mouse brain region to the human data (dorsolateral prefrontal cortex)^30^. We found no overlap of miRNAs that were regulated with age in the same direction in both data sets. An explanation for these results could be that the youngest age of 71 years in the human data roughly compares to the ending time point of the mouse data set with 28 months. Hence it is possible that the trends observed within the mouse data set over the entire life span could be present humans as well but cannot be captured during this short time course at the end of the human life span.

### Sex independent miRNA expression changes during aging

Apart from sex-specific miRNA expression changes, we recognized common signatures that we aim to verify in a joint analysis of all samples. Analyzing all samples together enables us to additionally investigate the pons. We calculated region-wise Spearman’s rank correlation coefficients of miRNA expression with age (Supplementary Fig. 4c, Supplementary Table 7) and aggregated the results resolved in unique and multiple miRNAs analogue to the sex-separated analysis (Supplementary Fig. 4d). We found less miRNAs significantly positively correlated with age in the combined analysis than in the sex-separated one and more miRNAs significantly anti-correlated. The higher number of significantly anti-correlated miRNAs was driven by 75 miRNAs in pons. We found 11 miRNAs significantly anti-correlated in multiple tissues (e.g. miR-18a-5p in olfactory bulb and pons) and 7 significantly positively correlated in multiple tissues (e.g. miR-155-5p in corpus callosum, mot. cortex, caudate putamen, medulla, choroid plexus, SVZ, thalamus and vis. cortex). Hence, we were able to detect more age-related miRNAs in the combined correlation analysis than in the sex-separated one due to higher sample size, including persisting top candidates, such as miR-146a-5p, miR-155-5p and miR-5100.

Comparing deregulated miRNAs within each region and each age stage, also revealed persisting aging signature in the combined analysis (Fig. 4a, Supplementary Table 8) as observed before in the sex-separated analysis (Fig. 3d). The number of miRNAs downregulated in pons varied from 46 miRNAs (15 months) to 397 miRNAs (18 months) (Fig. 4a). MiR-9-3p and eleven other miRNAs were downregulated in each comparison between 12 months to 28 months. Interestingly, we identified deregulations even though no significant anti-correlation was observed before in several regions (cerebellum, ent. cortex, hippocampus posterior, hypothalamus, choroid plexus). For example, in hippocampus posterior 196 miRNAs were commonly downregulated at 26 and 28 months of age even though no significantly anti-correlated miRNA was found. This indicates a strong expression decrease between 21 and 26 months which was not detectable via correlation. In choroid plexus, we determined 45, 122 and 179 significantly upregulated miRNAs at 15 months, 18 months and 21 months, respectively (Supplementary Fig. 4e). Seventeen of these were commonly deregulated in all three comparisons (including miR-155-5p and miR-146a-5p) and 103 were commonly deregulated at 18 and 21 months. In age stages later than 21 months no significantly upregulated miRNAs could be found. An aggregation of the significant results resolved in unique and multiple miRNAs analogously to the sex-separate analysis showed a high number of significantly upregulated miRNAs in choroid plexus (214 miRNAs) and significant downregulation in pons (94 miRNAs), SVZ (82 miRNAs) and choroid plexus (54 miRNAs) (Supplementary Fig. 4f). In general, we observed deregulation for all brain regions and age comparisons to 3 months for all expression levels (Supplementary Fig. 4g). In our previous study, which examined the expression by bulk sequencing, we observed peaks of deregulation at 12 and 18 months in the brain without any regional resolution^31^. The fact that we did not observe these peaks across all brain regions in this study highlights the importance of a region-specific investigation.

**Fig. 4:**
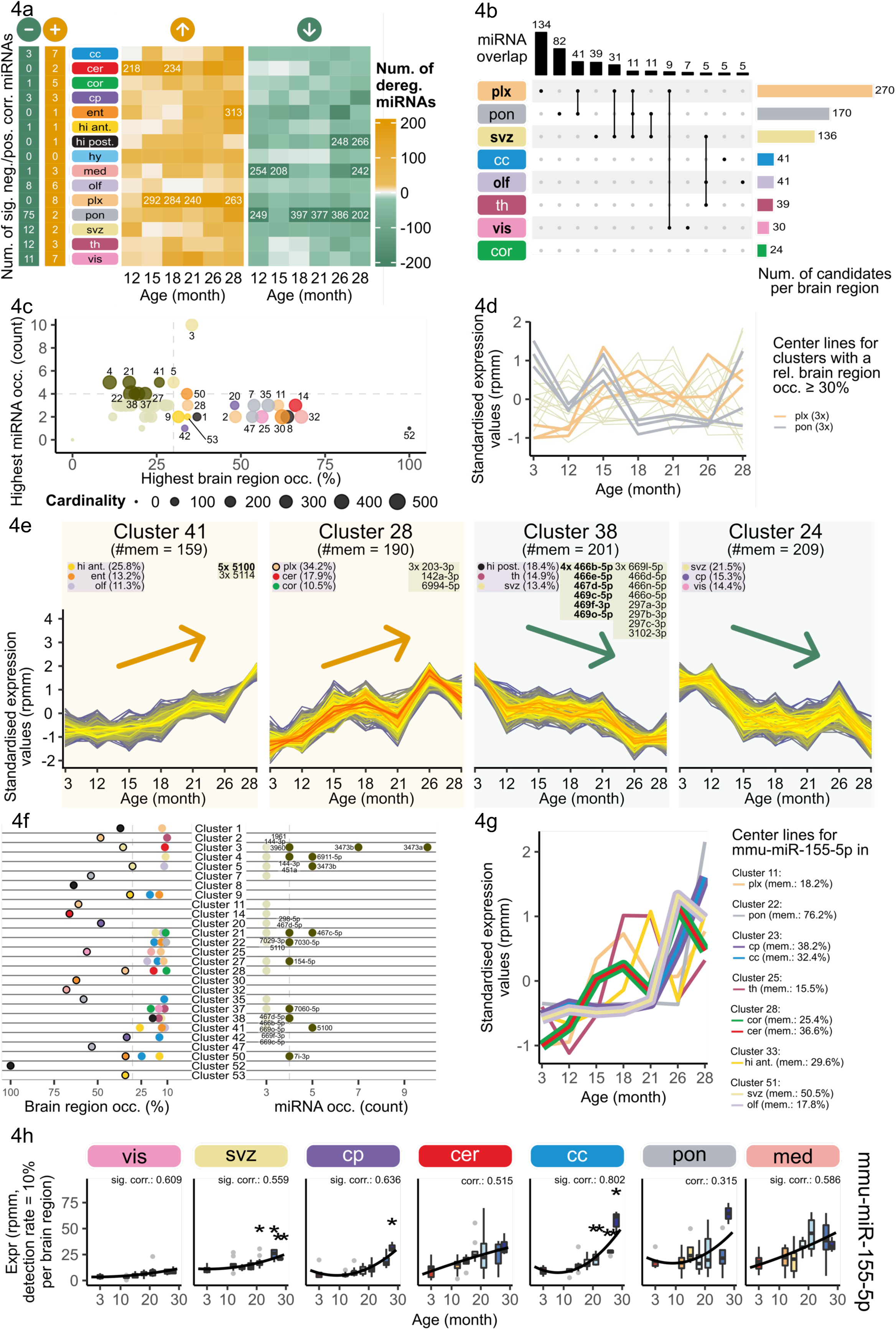
Global miRNA expression dynamics in brain aging across regions. In this figure, we consider all samples from our data set. **a** On the right heatmaps showing the number of deregulated miRNAs for each brain region regarding the comparison between each age stage to 3 months (fold change ≥ 1.5 or ≤ 1/1.5). Numbers in cell displayed when above threshold of 200 miRNAs. On the left numbers of significantly anti- or positively correlated miRNAs with age per brain region (using Spearman’s rank correlation coefficient with |R| ≥ 0.5, adjusted p-value < 0.05). **b** Upset plot providing an overview of miRNAs changing with age per brain region and presenting the uniqueness or overlap of these candidates. To determine these candidates, we determine the Spearman’s rank correlation coefficient for each miRNA in each brain region with age and filter for significantly positively or anti-correlated features (|R| ≥ 0.5, adjusted p-value < 0.05). Additionally, we perform a DE analysis for all comparisons between each age stage to 3 months. Additional candidates are those features for a brain region if the feature is significantly deregulated (abs. log2-fold change ≥ 1.5, adjusted p-value < 0.05) for at least one comparison. We highlight a brain region in bold if it is above the threshold in Fig. 2c concerning a high variance within age. For simplicity of the plot, we do not display any brain region with less than 10 candidates and combinations with less than 5 miRNAs. **c** Overview of the results of a trajectory clustering. We clustered the standardized time series for each miRNA and each brain region in k=53 clusters with a membership of at least 15%. Each dot corresponds to one cluster and the size of the dot corresponds to the number of trajectories in the cluster. We determine which brain region is most frequent in each cluster (relative to the cardinality of each cluster) and show the occurrence on the x-axis. If the value exceeds a threshold of 30%, we color that dot according to the brain region (see Fig. 1a). The y-axis shows the incidence of the most frequent feature in each cluster. If the value exceeds 4, we display the dot in a dark green color. Else it is colored in light green. **d** The displayed center lines belong to the clusters having a brain region occurrence higher than 30%, highlighted are the center lines for clusters with choroid plexus and pons. **e** Trajectories for four chosen clusters. Each line corresponds to the z-scored miRNA expression for one brain region. The two left plots show a steady increase with age and the two right plots show a steady decrease. At the top, we list the most frequent brain regions (rel. to the cardinality of that cluster) and the number of the most frequent miRNAs if the occurrence exceeds 10% in the brain region case and if the occurrence exceeds 3 in the feature case. Brain regions with a black border or bold miRNAs exceed the thresholds introduced in Fig. 4c. **f** A more detailed distribution of brain region occurrence (left) and miRNA frequency (right) for each cluster which exceeds any threshold from Fig. 4c. For simplicity, we do not show brain regions with an occurrence < 10% and a miRNA incidence < 3. For incidences exceeding the threshold from Fig. 4c (grey dashed lines), we provide the miRNA names and color the markers in dark green. For a better discriminability, we added a black border to the dots corresponding to the brain regions over the threshold. **g** The center lines of seven clusters containing trajectories corresponding to mmu-miR-155-5p. If trajectories for multiple brain regions were assigned to the same cluster and therefore have the same center line, we vary the line width for visualization purposes. **h** Boxplots showing the trajectories of miRNA mmu-miR-155-5p for seven brain regions. Asterisks highlight the significance of a deregulated comparison between that age and the control age (3 months). The Spearman’s rank correlation coefficient from the miRNA expression with age is displayed above each plot (significantly, if adjusted p-value < 0.05).

Age-related miRNAs from the deregulation or correlation analysis were determined analogously to the sex-separate analysis. We compared miRNAs identified as age-related within each brain region, evaluating unique and overlapping miRNA sets analog to the sex-separated analysis (Fig. 4b). There were many miRNAs uniquely associated with age in one region, especially 134 in the choroid plexus and 82 in pons. However, 41 common miRNAs were associated with aging in both these regions. The SVZ and choroid plexus also share 31 common aging miRNAs. SVZ, choroid plexus and pons shared 11 miRNAs, compromising the highest number of shared miRNAs and regions. The SVZ shared five age-related miRNAs with thalamus and the olfactory bulb, as the second highest amount of cross-region age-related miRNAs. To investigate the functionality of the identified age-related miRNAs, we compared the candidates per brain region to the previously published list of cholino-miRNAs^32^. Overlaps between these miRNAs regulating cholinergic genes and age-related miRNAs could indicate that the miRNA expression changes partially relate to altered acetylcholine signaling in aged individuals. We observed an overlap of seven age-related miRNA with the cholino-miRNAs in 7 out of the 15 brain regions (Supplementary Fig. 4h). In particular, mmu-miR-146a-5p stands out as it is significantly positively correlated in six brain regions and a previously known cholino-miRNA.

Further we examined whether the observed changes in miRNAs are driven by aging or can be related to mature miRNA decay or transcription process alterations. Per brain region, we examined a +/-10kb window around each significantly age-correlated miRNA on the same and the opposite strand and counted the number of significantly age-correlated miRNAs in the respective windows and accumulated the results across all brain regions (Supplementary Fig. 4i). As a reference, the average number of miRNAs in this defined neighborhood of a miRNA is 5.27 on the same strand and 0.25 miRNAs on the opposite strand. This analysis enabled us to explore whether the measured expression changes were accompanied by alterations on the opposite strand. No neighboring significant age-related miRNA was found in the analysis of the opposite strand. Additionally, we visualize the significantly correlated miRNAs with respect to their cumulative genomic coordinates (Supplementary Fig. 5a). Our findings indicate no significant enrichment on either the same or the opposite strand. This suggests that the observed miRNA changes are likely not a result of strand-specific transcriptional effects.

We performed a gene set enrichment analysis over the miRNAs ranked according to their significance and the direction of the Spearman rank correlation coefficients in all brain regions to explore the pathways targeted by the age-related miRNAs. This analysis revealed overlaps between six brain regions for the 50 most significant enriched and depleted pathways (Supplementary Fig. 6a). To name a few “GABA-ergic synapse” is depleted in hypothalamus, “Positive regulation of acetylcholine secretion, neurotransmission” in hypothalamus and thalamus and “Axon development” in pons. “Apoptotic process” is enriched in choroid plexus and vis. cortex and “DNA damage response” in choroid plexus. These results underscore the potential functional relevance of miRNA expression changes, as these are important pathways in the aging brain.

MiR-5100 as a cross-region age-related miRNA was observed in ten different brain regions (corpus callosum, cerebellum, ent. cortex, medulla, olfactory bulb, choroid plexus, pons, SVZ, thalamus and hippocampus anterior), miR-146a-5p in six (corpus callosum, mot. cortex, caudate putamen, olfactory bulb, choroid plexus and thalamus) and miR-155-5p in eight (corpus callosum, mot. cortex, caudate putamen, medulla, choroid plexus, SVZ, thalamus and vis. cortex). These observations indicated miR-155-5p, miR-146a-5p and miR-5100 as cross-sex brain aging miRNAs (Supplementary Table 9).

### Region-specific cluster organization of aging miRNA trajectories

To uncover continuous changes occurring over the entire measured lifespan other than correlations, we calculated miRNA aging trajectories for each miRNA in each brain region. Trajectory calculations were exclusively possible due to the continuous sampling within our data set. This analysis enabled us to study the time-related changes in greater detail than other aging studies using mostly a single young versus aged comparison. Subsequently, we clustered these trajectories in 53 clusters to identify common trajectories (Supplementary Fig. 7a, Supplementary Table 10).

Twenty clusters exhibited a strong region-specific signature as more than 30% of the trajectories in the cluster belong to the same brain region (Fig. 4c). Note, cluster 52 contained only one trajectory, hence we excluded it from further evaluations. Cluster 7, 35 and 47 revealed three distinct pons-specific aging miRNA trajectories dominated by a peak at 15 months and an overall decreasing expression (Fig. 4d). In contrast, we observed an overall increasing expression trend for the three choroid plexus region-specific clusters (cluster 2, 11 and 28). A loss of choroid plexus gene expression activity has been described in aging^33^, which could be due to increasing miRNA expression. Especially, cluster 28 and 41 are dominated by a steady expression increase within all miRNA trajectories (Fig. 4e). Although, cluster 41 exhibited an overall lower brain region occurrence. Therefore, we chose to investigate another characteristic within the clusters: the occurrence of a single miRNA trajectory originating from different regions (Fig. 4c). We found miR-5100 trajectories within the cluster 41 from five different brain regions (Fig. 4e; corpus callosum, cerebellum, ent. cortex, choroid plexus, thalamus). In the previous sex-separate analysis, we already identified this miRNA as age-related in both sexes with an expression increase. Cluster 28 contained miR-203 trajectories originating from three different regions (cerebellum, mot. cortex, choroid plexus). This miRNA negatively regulates NF-κβ signaling and microglia activation in neuronal injury^34,35^. It is therefore an interesting target in neuroinflammation regulation and brain injury, in which miR-203 inhibitors have already been successfully tested^36^. On the other hand, two clusters (24 and 38) exhibited a continuous decrease in expression. Neither of these clusters were region-specific. But cluster 38 contained trajectories from miR-466b-5p/466e-5p, 467d-5p and 469-families originating from four different regions, which were already identified as age-related in correlation and deregulation analyses.

We investigated the region-specificity of the clusters opposed to the occurrence of miRNA trajectories of the same miRNA originating from different regions in detail (Fig. 4c). If a cluster has a high region-specificity, there are no more than two to three miRNA trajectory occurrences in most cases. In contrast, clusters with multiple miRNA trajectory occurrences tend to have lower region-specificity. Out of the ten clusters (3, 4, 5, 21, 22, 27, 37, 38, 41, 50) harboring more than four times a single miRNA trajectory from different regions only three were deemed region-specific (3, 5, 50). So far, we only considered the most abundant miRNA or brain region per cluster. By investigating multiple occurrences of trajectories from the same miRNA and trajectories from the same brain region, we discovered that multiple clusters contained trajectories derived from different brain regions for the same miRNA and from the same brain region for different miRNAs. Clusters like 3, 4, 21, 22 and 38 contain multiple miRNA trajectories from at least four different brain regions (Fig. 4f). Especially in cluster 3, we observed that miR-3473a occurred ten times while the second most abundant mmu-miR-3473b occurred seven times. In cluster 38, five different miRNAs (466b-5p, 467d-5p, 669-family) occurred four times. In the case of brain regions, we never detected multiple brain regions above 30% within one cluster.

As mentioned before the cross-sex brain aging miRNA, miR-5100 was found multiple times in cluster 41. Apart from miR-5100, miR-155-5p and miR-146a-5p were significantly correlated with age in males and females and are known as key modulators of immune response^37^. We therefore investigated the clustering patterns of the three cross-sex brain aging miRNAs in detail. MiR-146a-5p trajectory center lines show an overall increase in age, especially clusters 22, 28, 41 and 51 (Supplementary Fig. 8a). Motivated by the cluster analysis, we take a closer look at the miR-146a-5p expression trajectories for all brain regions which show an overall increase for most brain regions (Supplementary Fig. 8b). Looking at cluster center lines containing trajectories from miR-5100, we observed an increasing tendence in particular for clusters 18, 22, 23, 30, 41 and 51 (Supplementary Fig. 8c). MiR-155-5p trajectories were observed in different clusters sharing an increasing center line (Fig. 4g). For example, the center lines of the clusters containing the miRNA trajectories for mot. cortex and cerebellum (28) increased towards 26 months. An increased expression most pronounced starting in 21 months was also visible in the clusters containing the miRNA trajectory for SVZ, olfactory bulb, caudate putamen and corpus callosum (51, 23). For the cluster containing the trajectory for pons (22), the increase was delayed to 26 months. As our initial correlation and deregulation analysis did not reveal the age-relation of miR-155-5p in pons and cerebellum, we chose to further investigate miR-155-5p expressions in all regions in detail as a promising cross-sex brain aging miRNA (Fig. 4h, Supplementary Fig. 8d). We observed an expression increase of miR-155-5p to different extents in all regions.

### Microglial age-driven miRNA expression changes

The results of the miRNA trajectory clustering revealed increasing expression tendencies for the three cross-sex brain aging miRNAs across all brain regions, even in regions previously not detected via correlation and deregulation analysis. This prompted us to investigate whether there could be a cell type driving the expression of these miRNAs. Mapping bulk miRNA expression back to cell types is a challenge within the field as there is no robust high-throughput single cell detection method for miRNAs since they are missing the polyA-tail most protocols depend on. However, miRNA expression patterns were previously measured using Cre recombinase-dependent miRNA tagging in motor neurons, all neurons, astrocytes and microglia^38^. MiR-155-5p and miR-146a-5p expression in the brain is likely driven by microglia, as both showed a prominent over 10-fold enrichment in microglia compared to brainstem^38,39^ (Fig. 5a, Supplementary Fig. 9a). Unfortunately, miR-5100 was not detected within this data set. To decipher whether the observed miRNA expression increase is driven by changes in cell type ratios or expression changes within the microglia, we sequenced microglia from young (3 months) and aged (21 months) mice collected via FACS sorting (Supplementary Fig. 9b, Supplementary Fig. 9c, Supplementary Table 1).

**Fig. 5:**
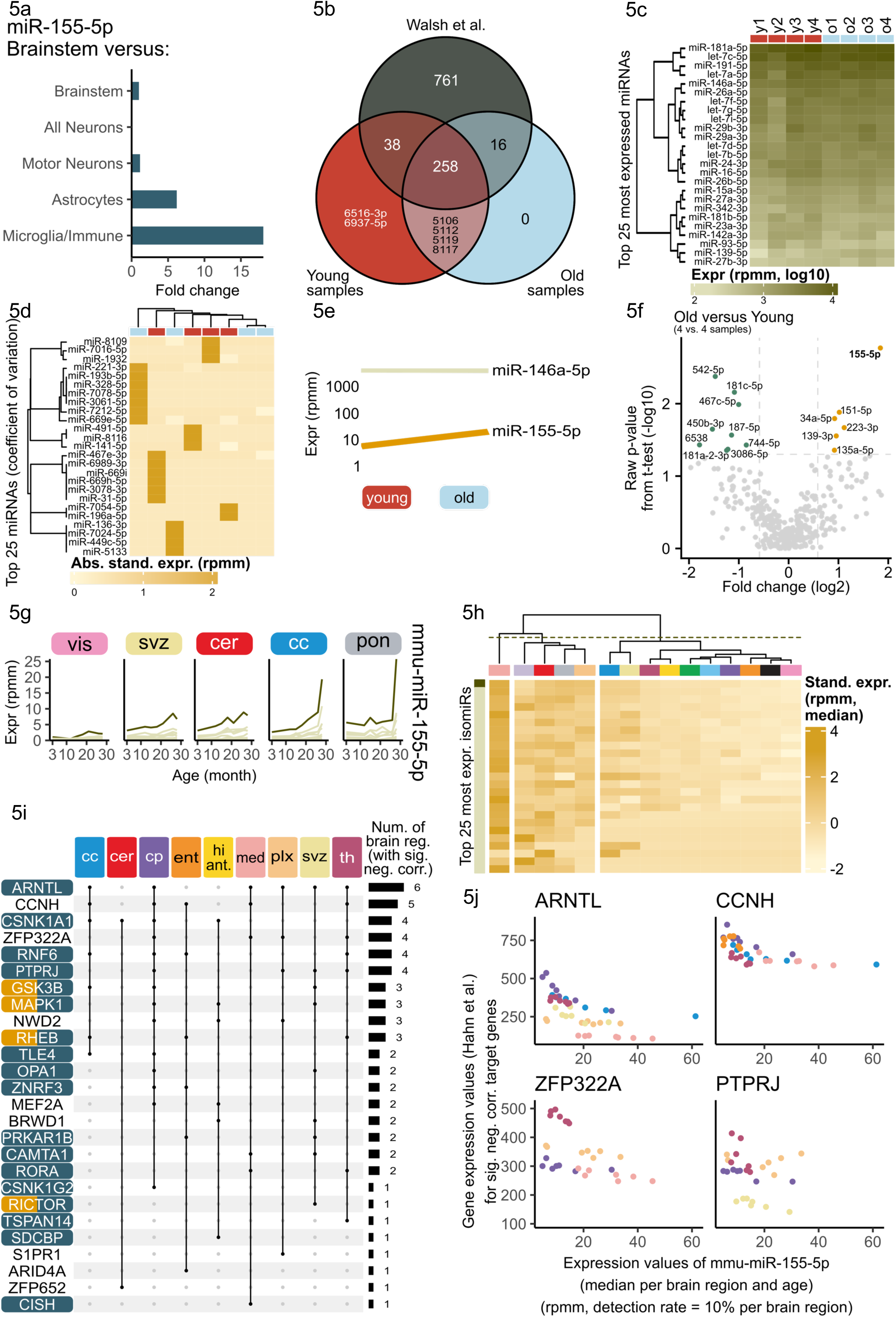
MiR-155-5p expression and target interactions in brain aging and microglia. **a** Adjusted from *CNS microRNA Profiles*^39^. We display the occurrences of mmu-miR-155-5p in different brain cell types via the fold changes between the cell type and the brainstem data. **b** Venn plot created in distinct mode for our microglia expression data derived from FACS sorted cells from young and aged mice compared to young microglia miRNA expression profiles derived from immunopanning^40^. **c** The top 25 most expressed miRNAs in our microglia data set (calculated over all samples) depicted in a log10-scale. The miRNAs are clustered using hierarchical clustering. **d** Top 25 miRNAs according to the coefficient of variation. We show the absolute standardized expression for each of the miRNAs and samples. Both the miRNAs and the samples are clustered via hierarchical clustering. **e** Line plot for two miRNAs showing the median values for young and old indicating the direction of deregulation and the expression. **f** Volcano plot presenting the DE analysis of the comparison old versus young showing the raw p-values. No miRNA is significantly deregulated considering the adjusted p-values. Nonetheless, we observe a strong upregulation for mmu-miR-155-5p marked in bold. Aging cohort: **g** Trajectories over the medians per age showing the expression of different isomiRs of mmu-miR-155 in five different brain regions. The canonical isomiR is highlighted as a dark line. **h** We calculate the 25 most expressed isomiRs decreasing from top to bottom over all samples. The averaged standardized expression values per brain region and isomiR are clustered according to the brain regions (using hierarchical clustering with complete linkage) and are split into three clusters. **i** Region-wise correlation values (Spearman’s rank correlation coefficient) for target genes (obtained from *miRTargetLink 2.0*^43^) of mmu-miR-155-5p using the mRNA data released by Hahn et al.^3^. The upset plot only shows the significantly anti-correlated genes ((|R| ≥ 0.3, adjusted p-value < 0.05) and the brain region in which the gene is significantly anti-correlated. Genes are colored by pathways: “mTOR Signaling Pathway” (yellow) and the “Regulation of cell communication” (blue). **j** The scatter plots are given for four of the significantly anti-correlated target genes shown in Fig. 5i. We see in each plot the brain regions in which this gene is significantly anti-correlated (indicated by the color) and the relation between the gene and the miR-155-5p broken down into median per brain region and age point.

First, we compared the miRNA expression of our young/aged microglia to data from Walsh et al.^40^ and found a strong overlap of expressed core miRNAs even though immuno-panning was used as isolation method (Fig. 5b). The most expressed miRNAs according to the median per samples were observed for young and aged, e.g. miR-181a-5p (Fig. 5c). Based on the coefficient of variation, we selected the top 25 miRNAs, as previously for which we clustered the samples. We found few miRNAs with rather distinct expression patterns in individual samples (Fig. 5d). Hence, we checked for the expression difference in our three brain aging miRNAs specifically. MiR-155-5p and miR-146a-5p were expressed in microglia above a threshold of 3 rpmm, miR-5100 was not expressed above this threshold. However, only miR-155-5p showed an increase in aged microglia (Fig. 5e). A detailed analysis of deregulated miRNA between these two groups revealed 59 miRNAs upregulated in aged microglia and 70 miRNAs downregulated (fold change greater or equal to 1.5 and smaller or equal to 1/1.5) (Fig. 5f, Supplementary Table 11). Within the upregulated miRNAs, miR-155-5p exhibited the highest fold change with a three-fold increase in aged over young (fold change: 3.58, raw p-value: 0.002 and adjusted p-value: 0.659). Hence, the increasing expression of miR-155-5p in several brain regions is likely driven by an increased microglial expression.

### MiR-155-5p as sex independent cross-region aging miRNA

Summarizing all our previous analysis, we considered miR-155-5p the most promising candidate for further investigation into its regulatory mechanisms. We examined whether its regulatory mechanisms contribute to known aging interventions such as dietary restrictions as well as young plasma injections, which were reported as beneficial for aged individuals^41,42^. Hence, we investigated the expression changes of miR-155-5p following 4 weeks of acute dietary restriction (aDR) and after young mouse plasma injections. We analyzed miRNA expression patterns in all previously collected regions of female mice aged 19 months, either fed *ad libitum* (AL) or treated with 4 weeks of aDR. And we analyzed samples from the young plasma injection cohort, which consisted of male 18 months old C57BL/6JN mice injected with young mouse plasma, following previously established protocols^42^. Filtering the samples analogously to the first dataset, we found an average read count per sample of over 21 million per group for the diet restriction experiment and 23 million for the plasma injection experiment, when aggregating all samples according to combinations of brain region and treatment (Supplementary Fig. 9d and 9e). On average, over 55% of the reads were mapped to miRNAs (Supplementary Fig. 9f and 9g) and 113 of 118 samples (96%) for the diet restriction experiment and 68 of 84 (81%) for the injection experiment passed our quality control (Supplementary Table 1). We included a total of 1345 and 1382 aligned miRNAs after an additional feature filtering for the diet restriction and injection experiment, respectively (cf. Methods). Across all regions, exclusively miR-451a showed a significant expression change during dietary restriction intervention (Supplementary Fig. 9h). An in-depth analysis of miR-155-5p expression in aDR and control did not reveal any persisting trends (Supplementary Fig. 9i). No significant expression changes were observed in any region after young plasma injections (Supplementary Fig. 9j). Similarly, for recurring young plasma injection, we did not detect any significant miR-155-5p expression changes compared to control groups, however we observed decreased expression in corpus callosum, cerebellum, caudate putamen, medulla, and choroid plexus and an increase in mot. cortex (Supplementary Fig. 9k). We hypothesize that the beneficial effects of these interventions are not crucially mediated via miRNA expression changes.

Sequence variations of mature miRNAs, so called isomiRs, occur naturally due to either altered cleaving patterns or sequence editing. IsomiR patterns can vary between different organs and in disease in differing ways to their archetype miRNA expression. IsomiRs can alter the target spectrum of a miRNA, hence we investigated whether the isomiR-archetype ratio is altered during the aging process. We found that isomiRs of miR-155-5p followed a similar trend of increase during aging in all regions, but the archetype form dominated over the isomiRs (Fig. 5g, Supplementary Fig. 10a). Furthermore, we checked whether there was a region-specific signature of isomiR expression for miR-155-5p. We discovered that in medulla miR-155-5p isomiRs showed a distinct expression pattern compared to all other brain regions (Fig. 5h).

As miR-155-5p increased during aging in all regions and this increase is likely driven by microglia, we investigated whether this increase leads to altered regulation of its mRNA targets. Therefore, we gathered all miR-155-5p targets using miRTargetLink 2.0^43^ (cf. Methods). Leveraging the matching bulk mRNA transcriptomic data from Hahn et al.^3^, we calculated the Spearman’s rank correlation coefficients between miR-155-5p and each target within each region. We denote a target gene significantly anti-correlated if the correlation value is smaller or equal than −0.3 and the adjusted p-value is smaller than 0.05. Eventually, twenty-six targets were significantly anti-correlated during aging in multiple regions (Fig. 5i).

Cyclin H (*CCNH*) and the basic helix-loop protein (ARNLT) were significantly anti-correlated in five and six different regions, respectively. Hence, increasing miR-155-5p expression in each of these regions lead to decreasing mRNA expression. The correlation strength in each region differs (Fig. 5j). In caudate putamen the miR-155-5p increase lead to a strong ARNTL decrease, whereas in medulla the mRNA decrease was less pronounced. We observed similar region dependent regulation strengths for ZFP322A (zinc finger protein) and PTPRJ (protein tyrosine phosphatase). PTPRJ activates MAPK1, which is an important player during extracellular signaling and various cellular processes. Additionally, we considered the functionally validated targets genes presented in Hart et. al^44^. Six of the 80 target genes showed a significant anti-correlation below –0.5 with mmu-miR-155-5p for at least one brain region, including the previously shown ARNTL (ADAM23, PCSK5, REPS2, NRCAM and NSG2, Supplementary Fig. 10b). In summary, we found multiple targets within important age-related pathways, such as the mTOR signaling pathway and targets within pathways of regulation of cell communication (Fig. 5i). Consequently, we focused on the additional transcripts of the mTOR pathway^45^ and their calculated Spearman’s rank correlations between the mRNAs and miR-155-5p (Supplementary Fig. 10c). For the brain regions corpus callosum, caudate putamen, ent. cortex, hippocampus anterior, choroid plexus, SVZ and thalamus a significantly negative correlation can be observed for multiple genes among these PDPK1, YWHAZ and CYCS. This highlights that a deregulation of miR-155-5p might not only affect its direct transcript targets but also other downstream targets within this pathway. Using the data from Keele et al.^46^, we explored the protein levels between young and aged mice in hippocampus, as we predicted that miR-155-5p expression increase leads to gene silencing by targeting the MEF2A transcript in hippocampus. In males, the MEF2A protein was less expressed in 18 months aged mice as compared to 8 months old mice (Supplementary Fig. 10d). We conclude that these analysis hint towards a regulatory effect of age-related microglial miR-155-5p expression changes on mTOR pathway gene expression.

## Discussion

In our aging brain region-specific miRNA expression atlas, we identified new region-specific miRNA expression signatures for multiple brain regions, especially in the olfactory bulb and cerebellum in males and medulla and pons in females. We were able to reproduce previous findings of miR-200 exclusively expressed in olfactory bulb in males and extend this knowledge to equal region-specific expression in females. This miRNA family was reported to play a crucial role in neurogenesis previously^22^. Furthermore, we found strong female region-specific expression patterns in medulla and pons that did not overlap with male expression patterns. Newly identified region-specific miRNAs from our analysis are therefore interesting targets to study region-specific functionalities.

In motor cortex and caudate putamen, we determined a strong sex-driven expression signature persistent over the entire lifespan. While analyzing the data in a sex-separated manner, we found that in females miRNA expression changes are most pronounced in choroid plexus, SVZ and hippocampus. While in males, expression changes in visual cortex, olfactory bulb and corpus callosum dominated.

The detailed analysis of miRNA expression patterns in brain regions during aging revealed sex-independent miRNAs increasing with age in multiple brain regions. We identified miR-146a, miR-155-5p and miR-5100 via correlation, deregulation and cluster analysis as cross-sex brain aging miRNAs. Especially miR-155-5p, which we identified as a global aging miRNA before^31^, is an interesting target for further study in the aging brain context. This miRNA has been shown to regulate leukocyte adhesion at the inflamed BBB^47^. Additionally, miR-155-5p has been associated with neurological disorders such as Alzheimer’s^10^ and the murine model of multiple sclerosis^48^, making it a promising therapeutic target. We detected an increase of miR-155-5p expression in aged microglia in our study. Increased secretion of miR-155-5p from microglia mediates inflammatory neuronal cell death and therefore plays a pro-inflammatory role^49^. Whether the increased microglial expression of miR-155-5p in aged individuals is limited within this cell type or is actively secreted into the microenvironment remains to be studied. Secretion of miR-155-5p via e.g. exosomes could contribute to the “inflammaging” phenotype, as miR-155 has been proposed as a central regulator in CNS-related inflammation by affecting microglia^50^. In the disease context (AD), miR-155 together with interferon-γ signaling mediates a protective microglial state^10^. A microglia specific deletion of miR-155 reduced amyloid-β pathology in AD mouse models but caused hyperexcitability and seizures^51^. The distinct isomiR expression in certain brain regions, like medulla for miR-155-5p calls for further investigation. An altered target spectrum is a potential functional consequence of strong isomiR expression. In our data set, we discovered altered isomiR expression for miR-155-5p and an expression change in a target gene (CISH) exclusively in medulla. Additional studies could use this dataset together with functional validation experiments to advance understanding of isomiR regulatory properties in general and specifically the aging context.

Limitations of our study include in addition to restricted transferability of all findings to humans due to usage of an inbred mouse line and the sparsity of human tissue to create a comprehensive time course, the limitations arising from technical challenges of analysis of cell-type specific miRNA expression patterns. Commonly used single-cell sequencing techniques cannot be applied to assess mature miRNA expression due to the lack of polyA-tails, which are essential to most protocols. Hence, we had to resort to FACS to sort for microglia. Selecting antibodies used for the sorting process requires a careful balance to specifically select the cells of interest. In our strategy we chose to exclude CD45+ cells to distinguish resident microglia from infiltrating immune cells. However, thereby we exclude a subset of resident microglia, as the expression of CD45 can also increase with age in resident microglia. Another limitation is the question whether different microglial states in dietary restriction and young plasma injection exist with respectively specific miRNA expression patterns. Changes in cell type composition and especially cell type states could drastically influence miRNA bulk expression patterns. These limitations further underline the need for detailed studies of cell-type specific miRNA expression patterns and even different cell-type states. Due to limited access to aged mice this was not possible in this study. Especially acquiring samples from aged female mice is challenging, therefore our experimental microglia data is limited to male samples. In addition to human data, validation of brain aging miRNAs in other mouse strains would strengthen our hypotheses as differing miRNA expression patterns in different brain regions were previously reported^52^.

In conclusion, we identified microglial derived miR-155-5p as the most interesting therapeutic target. As it regulates broadly cellular communication pathways during aging and increased sex-independent manner across all brain regions. MiR-146a-5p is another interesting candidate, because we also observed it’s sex-independent cross-region expression increase in this study. In addition to these cross-region targets, we identified unique aging signatures in e.g. choroid plexus, subventricular zone and pons as interesting targets to study in context of functionality of these brain regions. The samples size of this study enabled us to even identify sex-specific aging signatures that are currently heavily understudied. Future studies with more human samples are called for to validate these findings in humans to enable translation into clinical applications. In sum, this atlas offers all researchers a comprehensive map of miRNA expression across different brain region in a sex-specific manner, which was previously nonexistent with possibility to also go in-depth into isomiR analysis, including the multiple time points to study aging signatures. The study is a powerful resource as a reference, especially for age-related and sex-specific miRNA expression changes across the entire life span in mice.

## Author contributions

Conceptualization: A.K., T.W-C., O.H., V.W.; Methodology: N.L., V.W.; *in-*vivo: microglia experiments: N.L, I.G., A.S., V.W. Library Preparation: N.L, A.B., V.W.; Software: A.E.; Resources: O.H., T.W-C.; Data Curation: A.E., V.W.; Writing – Original Draft: A.E., V.W.; Writing – review & Editing: O.H., U.F., A.K., T.W-C.; Visualization: A.E., A.K., V.W.; Supervision: E.M., A.K., T.W-C.; Funding Acquisition: A.K., T.W-C., E.M.

## Data and code availability

All sequencing data are available for download from NCBI’s Gene Expression Omnibus with the accession numbers GSE282205 and GSE282207 for the brain aging, invention experiments and the microglia data, respectively. The data will be made public at final publication. We built a web service that offers interactive access to processed bulk-sequencing data from the brain aging cohort, as well as experiments on dietary restriction, young plasma injection, and microglia. The web service is accessible via https://ccb-compute2.cs.uni-saarland.de/brainmirmap. Human ROSMAP data is available via Synapse^53^. The code supporting the analysis within this paper is available at GitHub at https://github.com/CCB-SB/2024_brain_aging_miRNA.

## Acknowledgements

We thank all members of the Wyss-Coray and Meese lab, as well as all members of the Keller lab for feedback and support. This study is funded by the Saarland University (A.K.), the DAAD (V.W.), NIH Pathway to Independence Award 1K99AG088304-01 (I.H.G), AHA-Allen Brain Health and Cognitive Impairment Cross-Network Collaborative Grants (23BHCICG1188316; N.L.) and the M.J. Fox Foundation (MJFF-021418; A.K. & T.W-C.). Computational resources used within this study were financed through the DFG project 466168625 (A.K.). N.L. is a MAC3 Dementia and Ageing Fellow supported by MAC3 Impact Philanthropies. Special thanks to Phillip Gross (Georgetown University Medical Center) and Ruben Garcia Martin (Centro Nacional de Biotecnología CSIC (CNB), Madrid) for their support.

## Methods

### Samples

As previously described male and female C57BL/6JN mice from the National Institute of Aging colony (Charles River) were shipped to the Stanford ChEM-H animal facility (Palo Alto), where they were housed at least one month before euthanasia^3^. For each age group of 3, 12, 15, 18 and 21 months 5 female mice and 5-6 male mice were used; age groups 26 and 28 months consisted only of 5 and 3 male mice respectively. Animals were housed in cages of 2-3 mice, with a 12/12 light/dark cycle, at 19.4 – 22.8 °C and provided with food and water ad libitum. Over the course of four days the sample collection was performed between 10 am and 12 pm. Mice were anaesthetized with 2.5% v/v Avertin, 700 µl of blood was drawn via cardiac puncture and followed by transcardial perfusion with 20 ml cold PBS. After immediate removal of the brains, the organs were snap-frozen by submission in liquid nitrogen-cooled isopentane (60 seconds) and ultimately stored at −80°C before further processing. The respective regions were dissected via slicing and atlas-guided tissue punching while frozen. Using a metal brain matrix coronal sections of 1 mm thickness were sliced with .22 razor blades (Ted Pella, 15045; VWR, 55411-050). Regions of interest (1.5mm and 2mm diameter) were dissected quickly from the right hemisphere of these sections using disposable biopsy punches (Alimed, 98PUN6-2, 98PUN6-3). The following 15 regions were collected: three cortical regions (motor cortex, visual cortex and entorhinal cortex), anterior (dorsal) and posterior (ventral) hippocampus, hypothalamus, thalamus, caudate putamen (part of the striatum), pons, medulla, cerebellum and the olfactory bulb, corpus callosum, choroid plexus and the subventricular zone. Four regions were collected in the following order, as the collection required overlapping punches: (1) motor cortex, (2) caudate putamen, (3) subventricular zone, (4) corpus callosum. All animal care and procedures complied with the Animal Welfare Act and were in accordance with institutional guidelines and approved by the institutional administrative panel of laboratory animal care at Stanford University. RNA was isolated using the RNeasy 96 kit (Qiagen, 74181) and a TissueLyser II (Qiagen, 85300), according to RNeasy 96 Handbook protocol “Purification of Total RNA from Animal Tissues using Spin Technology” without the optional on-plate DNase digestion.

### Aging interventions

Young Mouse Plasma (YMP) was collected following the protocol described by Villeda et al.^42^ Briefly, C57Bl/6J male mice aged 2 months were anesthetized with 2.5% v/v Avertin after being group-housed. Around 700 μl of blood was drawn via cardiac puncture prior to transcardial perfusion. 15 μl of 250 mM EDTA (Thermo Fisher Scientific, 15575020) were used to collect blood. The mixture was centrifuged at 4°C for 15 minutes at 1,000g to obtain plasma. The plasma from 20-25 mice was pooled together and dialyzed in 1X PBS using cassettes (Slide-A-Lyzer Dialysis Cassettes, 3.5 kDa molecular weight cut-off, 3-12 ml) before being frozen at −80°C. For plasma transfer experiments, 18 months old male C57BL/6JN mice were injected retro-orbitally with 150 μl of YMP per injection. Prior to injection, mice were habituated by being placed on the procedure table in their cage. Injections were administered every 3-4 days, alternating between the left and right eye to allow for recovery. Mice were rested for four days before tissue collection.

For the aDR study with C57BL/6JN mice, 18-months-old mice were randomly assigned to AL or aDR. aDR treatment was initiated by transferring mice from AL to 10% aDR for 7 days. After that, aDR was increased to 25%. aDR animals were fed once per day between 3-5 p.m., and all animals were checked daily for their well-being and any deaths. For the first 16 days, weights were checked daily. Mice were euthanized at the ages of 19 months. All mice were euthanized in the morning within a period of 6 hours prior to the regular feeding time of the DR mice.

The aDR study with C3B6F1 mice was performed in accordance with the recommendations and guidelines of the Federation of the European Laboratory Animal Science Association (FELASA), with all protocols approved by the Landesamt für Natur, Umwelt und Verbraucherschutz, Nordrhein-Westfalen, Germany (84-02.04.2015.A437). Female F1 hybrid mice (C3B6F1) were generated in-house by crossing C3H/HeOuJ females with C57BL/6NCrl males (strain codes 626 and 027, respectively, Charles River Laboratories). Five female animals were housed as a group in individually ventilated cages under specific-pathogen-free conditions with constant temperature (21 °C), 50–60% humidity and a 12 h/12 h light/dark cycle. For environmental enrichment, mice had constant access to nesting material and chew sticks. All mice received commercially available rodent chow (ssniff R/M-Low phytoestrogen, ssniff Spezialdiäten, Germany) and were provided with filtered water ad libitum. aDR animals received 60% of the food amount consumed by AL animals. aDR treatment was initiated at 20 months of age by directly transferring mice from AL to 40% DR. aDR animals were fed once per day, and all animals were checked daily for their well-being and any deaths. Mice were euthanized at the ages of 24 months. All mice were euthanized in the morning within a period of 3 hours prior to the regular feeding time of the DR mice. Mice were euthanized by cervical dislocation, and tissues were rapidly collected and snap-frozen in liquid nitrogen.

The cohort of mice treated with YMP or PBS were housed at the Palo Alto VA animal facility under a 12 h/12 h light/dark cycle at 68–73 °F under 40–60% humidity. All experiments were performed in accordance with institutional guidelines approved by the VA Palo Alto Committee on Animal Research. Euthanasia and organ collection was conducted in the same way as the aging cohorts.

### Microglia isolation

Microglia from young and aged mice (3 and 21 months, C57BL/6, males) were isolated via FACS-sorting following the below described protocol. In brief, mice were anaesthetized with Avertin and perfused with 20 mL ice cold DPBS. Brains were dissected, hemispheres separated, and olfactory bulb and cerebellum removed. Single-cell solutions were created for each hemisphere via mincing and douncing the tissue, the solution was filtered (70µm cell strainers, Falcon 352350) and finally centrifuged (400 x *g*, 10 mins, 4°C). After resuspension in MACS buffer and addition of myelin removal beads (Miltenyi Biotech, 130-096-433), solutions for each hemisphere were loaded on LD columns (Miltenyi Biotech, 130-042-901) to remove myelin. Columns were washed twice with MACS buffer. Cells were pelleted via centrifugation (400 x *g*, 5 mins, 4°C) and resuspended in FACS buffer. FC blocking antibody was added and incubated for 5 mins. Primary antibodies (CD11b-FITC (Biolegend 101206); CD-45-BUV396 (Biolegend 50-162-785) and CD206-APC (Biolegend 141707)) were added and incubated for 30 mins on ice. After another centrifugation, samples were resuspended in 0.5 mL FACS buffer and Sytox Blue (Thermo, S34857) was added for live cell labelling. Cells were sorted on a MA900 Multi-Application Cell Sorter (Sony Biotechnology) and selected by gating for live single cells (FSC-A/SSC-A). Further, cells were gated for CD11b^+^; CD45^low^; CD206^-^ and sorted directly into 1.5mL tubes containing 50µl FACS buffer. As CD11b^+^/CD45^+^ brain cells are mainly infiltrating macrophages (PMID: 32381088) we exclude these macrophages with our sorting strategy. FACS-isolated CD11b^+^/CD45^low^/CD206^-^ brain myeloid cells are referred to as microglia in this manuscript. Between 10,000 and 40,000 cells per sample were collected. For aged mouse sample 4, two mice were pooled together to reach the threshold of minimal 10,000 cells as input for RNA isolation. After collection was completed 1 mL QIAzol was added instantly. After vortexing, samples were stored at −80°C until RNA isolation. Since cell sorting provides little input material for RNA isolation, the standard miRNeasy microKit (QIAGEN, cat. no. 217084) was used with standard protocol for elution of separated fractions for RNAs above and below 200 nt, to concentrate the miRNA input for library preparation. Library preparation of microglia miRNAs was optimized and performed as described below, except for the RT-primer input, which was altered to a 1:5 dilution and amplification PCR was 25 cycles long.

### Library Preparation

The MGIEasy Small RNA Library Prep Kit (Item 940-000196-00) was used for library preparation on the high-throughput MGI SP-960 sample prep system according to the manufacturer’s protocol. In principle this library preparation method works by ligating 3’- and 5’-adapters to all RNAs in each sample. During reverse transcription (RT), specific RT primers that bind to the adapters are used to generate cDNA and introduce sample specific barcodes. Amplification of this cDNA is performed via a 21-cycled PCR. Size-selection of this PCR product is performed via magnetic beads (AMPure Beads XP, Beckman Coulter). To focus on the small RNAs a size of around 110 bp was selected, this was checked using an Agilent DNA 1000 Kit (Agilent Technologies). The concentration of each sample was measured by a QuBit 1x dsDNA High Sensitivity Assay (Thermo Fisher Scientific). Each library in this dietstudy consisted of 16 samples, barcoded with the following barcodes: 1–4, 13–16 and 25–32. All samples of one library were pooled after concentration measurement in an equimolar fashion to reach a concentration of 4.56 ng µl-1 for each sample in each pooled library. After circularization the pooled libraries were sent for sequencing.

### Sequencing & Data Analysis

Samples were single-end sequenced on the BGISEQ500RS using the High-throughput Sequencing Set (SE50) (Small RNA) as a service provided by BGI, Hong Kong. We utilized miRMaster 2.0^55^ with standard settings on the given data sets which performed an alignment against the mouse genome (GRCm38) and a mapping against miRNAs using miRBase^56^ (version 22.1) using Bowtie^57^ (version 1.2.3) with the options “-m 100 —best —strata” to obtain the raw counts for miRNAs and their isomiRs, for lncRNAs, piRNAs, rRNAs, scaRNAs, snoRNAs, snRNAs and tRNAs. The miRMaster pipeline denotes any fragment as either tRNA or lncRNA even though it cannot detect them at full length as we are size selecting for miRNAs as our RNAs of interest^58^. Additionally, we gathered the alignment and mapping information which were produced during the run. For the further analysis we worked with a misclassification rate of 1. We normalized the raw counts with a rpmm-normalization to be able to ensure comparability between the different samples. Next, we filtered the samples and features. We kept only the samples for which more than 2 million reads could be aligned to the mouse genome. Feature filtering was performed by checking if for at least 10% of the samples for at least one group the raw count exceeded or was equal to 5. Subsequently, for the aging cohort, we obtained 844 sequenced samples and 1966 miRNAs (9139 lncRNAs, 30930 piRNAs, 356 rRNAs, 51 scaRNAs, 1542 snoRNAs, 1390 snRNAs and 408 tRNAs) from which we kept 828 samples and 1174 miRNAs (3424 lncRNAs, 605 piRNAs, 194 rRNAs, 27 scaRNAs, 672 snoRNAs, 705 snRNAs and 404 tRNAs). For the diet restriction and injection experiment, we added a further quality control step. If we were left with three or less samples per brain region after the sample filtering, we additionally discarded all samples of this brain region. This resulted in 118 sequenced and 113 kept samples for the diet restriction experiment and 68 out of 84 samples for the injection experiment. For both, we mapped 1966 features and obtained 1345 and 1382 features after filtering, respectively. The microglia data set consisted of 8 samples of which all samples and 419 features survived the filtering using an adjusted threshold of 1.8 million reads. All isomiR expressions were also normalized by rpmm-normalization and included the same samples as their corresponding expression tables and the isomiR forms connected to these miRNAs. Therefore, we did not perform a separate filtering for the isomiRs. We called a feature most expressed if its median value over all samples was the highest compared to the other miRNAs. We denoted a feature as expressed in a brain region if at least 10% of the samples of that brain region had a raw count higher or equal than 5. Human data from ROSMAP^29^ was processed with miRMaster 2.0^55^ with default settings. We performed the same pre-processing as explained above up to the feature and samples filtering where we changed the thresholds for the number of aligned reads to 10,000 and the raw count value to 2 what need to be fulfilled for 10% of the samples for either the male or female samples. So, after filtering we obtained 203 samples and 280 miRNAs.

We use all features for the Uniform Manifold Approximation and Projection (UMAP) calculation which was done according to McInnes et al.^59^ and was colored by information from the metadata. The parameters were selected visually from the results of a pool of linear combinations of default parameters. We prepared the expression matrix for the calculation by standardizing it (z-scoring). As parameters we chose 0.25 as the minimum distance, as metric we used the Euclidean and we started with a random initialization. For the young samples, we used 5 as the size of the local neighbourhood and for the samples from all ages we used a local neighbourhood of 10.

We calculated per brain region the expressed features (raw count greater or equal than 5 for at least 10% of the brain region’s samples) for every of the seven RNA classes. From those, we determine relative values to the total number of raw counts of all expressed features from all RNA classes of each brain region individually. We display these values in a stacked bar plot per brain region colored by the RNA classes. Due to the different raw count distributions of the RNA classes, the share of miRNA becomes less prominent compared to the mapping results. By the same approach, we obtain relative values of the RNA classes per brain region on an age-resolved level to gain an overview of the raw count composition over time. Per brain region we show the relative data points as a scatter plot for each age colored again by the RNA classes. A third-degree polynomial fitted per RNA class to the respective data points indicates the age-trend of the raw count composition. Using the rpmm-normalized and filtered expression of each RNA class, we calculate the correlations with age of each feature per brain region using Spearman’s rank correlation coefficients. Showing the resulted correlation values in a ridgeline plot (like a histogram) joint over the brain regions per RNA class illustrates in which RNA class we obtain the most positive and negative correlations with age.

Box plots for features per brain region resolved by age were calculated. The box borders correspond to the 25th (Q_!_) and 75th Percentile (*Q*_3_), the middle line to the median and whiskers to the minimum (maximum) of the minimum value or *Q*_!_ − 1.5 · IQR (maximum value or the *Q*_1_ + 1.5 · IQR) where IQR determines the interquartile range. Solid grey dots in the plot indicate the potential outliers in the data. We fitted a polynomial regression line with degree 2 to the data. For male and female separately, we calculated the Spearman’s rank correlation coefficients for each feature with the age per brain region. The corresponding p-values were adjusted using the Benjamini-Hochberg procedure.

To quantify the difference of a brain region to the average brain, we used the coefficient of variation given by the ratio of standard deviation and mean value per feature. We selected the 50 miRNAs yielding the highest value. To differentiate features from the brain average for a particular brain region, we calculated per brain region the median expressions and standardized each feature (z-score). Thus, we called a feature different from the brain average for a specific brain region if the absolute z-score was higher or equal than 0.5. For the so obtained binarized table, we applied a hierarchical clustering using complete linkage to cluster the binarized expression profiles into four clusters.

Next, for the analysis regarding the sex comparison Male vs Female, we discarded all samples from the brain region pons due to a lack of samples for older ages in the female case. All Principal Variance Component Analysis^60^ (PVCA) in this publication were processed with properties given by the metadata all of which could be seen in the PVCA results presented as bar plots together with two-way interaction terms (containing all combinations of the properties). For clarity, only two-way interaction terms with a non-vanishing observed variance were shown. The residual contains the observed variance which was not covered by the used metadata. While a PVCA is normally used to reveal batch effects within the data, we used it to identify properties suggesting a high impact in the dataset guiding a further analysis. In a second investigation, we applied the PVCA to every brain region individually for the properties age and sex. We applied a DE analysis per brain region for the sex comparison Male vs Female. We calculated the fold changes for each feature by dividing the geometric median over all male samples by the geometric median over all female samples in every brain region. For any further analysis we removed the features exhibiting no deregulation (fold change of 1). To obtain the adjusted p-values we performed a Welch’s *t*-test which is an adaption of Student’s *t*-test (for simplicity we always write Student’s *t*-test instead of Welch’s *t*-test) with the corresponding samples and adjusted the obtained p-values using the Benjamini-Hochberg procedure. We called a feature significantly up- or downregulated if the adjusted p-value was smaller than 0.05 and in case of an upregulation (down-) if the fold change was greater or equal than 1.5 (smaller or equal than 1/1.5). We performed a gene set enrichment analysis for all features. Therefore, we ranked the features with a positive (negative) log2-transformed fold change increasingly (decreasingly) according to their significance (p-value). Subsequently, the enrichment analysis was performed using miEAA 2023^54^ with the combined list of ranked features.

We called a feature significantly positively or negatively correlated with the age if the adjusted p-value was smaller than 0.05 and positively (negatively) correlated if the correlation value was greater than 0.5 (smaller than −0.5). Additionally, we denoted a feature as “unique” if it exhibited a significantly positively or significantly negatively correlation within only one brain region. If this held for more than one brain region for the same direction, we called it “multiple”. Subsequently, we determined the deregulated features per brain region by calculating the fold changes analog to above for the comparisons of every older age (12m, 15m, 18m, 21m and in the case of male add. 26m, 28m) versus the control age of 3m again for male and female separately. Therefore, we used the medians of each group. Again, we determined the p-values with the Student’s *t*-test and adjusted them by using the Benjamini-Hochberg procedure. Analogously to above for the correlation approach, we introduced the property “unique” or “multiple” for features which were significantly up- or downregulated in at least one age comparison within a brain region. If this feature was in only one brain region significantly deregulated, we called it “unique” and for more than one, we called it “multiple”. To close the side-by-side analysis, we combined the male and female data back to one dataset for which we calculated the unique and multiple features for the two approaches like explained above. We call a feature a candidate if it was significantly positively or negatively correlated with age in at least one brain region or significantly up- or downregulated in at least one age comparison for a brain region. If the direction changed between the brain regions the features just counted as one candidate.

The human miRNA data from ROSMAP^29^ was binned for the age of death in three half-open intervals [71,81), [81, 92) and [92, 103) in years. Afterwards, we perform a DE analysis between the oldest and the youngest group using the Student’s *t*-test and Benjamini-Hochberg to adjust the p-values for multiple testing. Again, the fold changes were calculated with the geometric means. Additionally, we determined Cohen’s d for every feature to obtain a measure for the effect size^61^.

Madrer and Soreq^32^ introduced a set of miRNAs which are predicted to target cholinergic genes. We intersect this list of mIRNAs with the miRNAs of our dataset found to be significantly positively or negatively correlated with age individually for each brain region.

For each significantly age-correlated miRNA, we investigated a 10kb range on the same and the opposite strand and determined significantly age-correlated miRNAs. Hence, we obtained a tuple of the number of feature candidates on the same and the opposite strand. Summing up the occurrences of this tuples for each significantly age-correlated miRNA within each brain region yields a global overview. Per brain region, we conduct a gene set enrichment analysis for all features. Therefore, we ranked the features with a positive (negative) Spearman rank correlation coefficient with age increasingly (decreasingly) according to their significance (p-value of the correlation). We used miEAA 2023^54^ with the combined list of ranked features to obtain the enrichment analysis results.

Using a c-means clustering^62^, we clustered the standardized trajectories given by the miRNAs resolved in the brain regions (total: 15 brain regions × 1174 features = 17610 trajectories). We determined the number of used clusters by visually inspecting the minimum centroid measurements for all cluster numbers from 2 to 200. From the method, we obtained a percentage value for each trajectory and each cluster containing the probability that the trajectory belongs to this cluster. Using this measure, we assigned each trajectory to the cluster where it exhibits the highest membership and afterwards discarded the ones with a membership lower than 15%. Therefore, each cluster with its specific aging trajectory has a unique composition of tuples of miRNAs and brain regions. Each tuple only occurs once across all clusters. We called a cluster brain region specific if more or equal than 30% of the elements in this cluster belonged to only one brain region and feature specific if 4 or more occurrences of one feature could be found in the cluster.

miRTargetLink 2.0^43^ provided 82 target genes for the feature mmu-miR-155-5p considering all functional ones. Using the gene data from Hahn et al.^3^, we obtained for 66 of the target genes expression values from 809 samples matching to the miRNA samples. We calculated the Spearman’s rank correlation coefficients between the target genes and the miRNA 155-5p for every brain region and the corresponding adjusted p-values with the Benjamini-Hochberg procedure. For this part of the analysis, we called a gene significantly negatively correlated if the correlation value was smaller or equal than 0.3 and the adjusted p-value was smaller than 0.05. Further, we only investigated target genes, and their expression values resolved in the ages for only the brain region for which they were significantly negatively correlated with mmu-miR-155-5p. In the mTOR pathway are 67 genes included^45^. For 63 genes we calculated the Spearman rank correlation coefficient using the mRNA data presented by Hahn et al.^3^. As an adjustment method we used the Benjamini-Hochberg procedure. Analog to above we calculated the Spearman rank correlation coefficients for the miRNA mmu-miR- 155-5p and 66 from the 80 functionally validated target genes introduced by Hart et al.^44^ for each brain region. We consider genes for which at least one brain region exhibit a significant (adjusted p-value < 0.05, Benjamini-Hochberg procedure) correlation value below −0.5.

For data introduced in Keele et al.^46^ and published via their web interface “Aging B6 Proteomics”, we show the intensities for MEF2A in hippocamous as box plots. The box plots are build like above due to the low number of samples per box we added black cirlces visuallising the exact data points within the plots.

The topmost expressed isomiRs were calculated analogue to the most expressed miRNAs. The brain region clustering for the standardized isomiR expression data was achieved by hierarchical clustering using complete linkage. The features were ranked based on their expression level.

A visualization of the data set from Hoye et el.^38^ was implemented accordingly to previous publications^39^. The top miRNAs in the microglia dataset were obtained analogue to above. The clustering of the features was done by hierarchical clustering using complete linkage. Analogously to the aging cohort, we calculated the top features based on the coefficient of variation and clustered the standardized expression values by samples and features (hierarchical clustering using complete linkage). A DE analysis within the microglia data was done analogue as before for the comparison Old versus Young. We selected 5 upregulated (greater or equal to 1.5) and 5 downregulated features (smaller or equal to 1/1.5) by their adjusted significance for further analysis. For the diet restriction and injection data set, we obtained a DE analysis analogue to above for the comparison Treatment versus Control and old mice with young plasma versus PBS for all samples and split in the different brain regions. We calculated violin plots for features per brain region resolved by treatment type. The violin displays the density of the data points. Grey dots in the plot represent all data points used for this violin.

The analysis and figures were produced via snakemake pipelines^63^ (version 7.18.2) using R (version 4.2.2) and Python (version 3.11.0). Data analysis was done using the packages data.table (version 1.14.6), ggrepel (version 0.9.2), reshape2 (version 1.4.4), stringr (version 1.4.1), mfuzz^64^ (version 2.58.0) and deseq2^65^ (version 1.38.0). The umap calculation in Python was implemented using the package umap-learn^59^ (version 0.3.10), numpy (version 1.19) and pandas (version 0.24.2). The heatmaps were created with ComplexHeatmap^66^ (version 2.14.0) and circlize (version 0.4.16) all other figures with ggplot2 (version 3.3), gridtext (version 0.1.5) and fontawesome (version 0.4.0).

## Supplementary information

**Supplementary Fig. 1: a** Overview of the aligned (green) and not aligned (grey) reads against the mouse genome. Combined per brain region and age. Values are given in millions. **b** Reads mapped to RNA types per sample. Brain regions are highlighted at the left side. **c** UMAP for all regions and all features (miRNAs, lncRNAs, piRNAs, rRNAs, scaRNAs, snoRNAs, snRNAs and tRNAs) colored by brain regions as indicated in Fig. 1a. **d** UMAP of all samples and all features colored by sex. Analogue to Supplementary Fig. 1c. **e** UMAP of all samples and features colored by age. Analogue to Supplementary Fig. 1c. **f** For each brain region and RNA class, we determined the composition of expressed RNA counts, considering only RNAs with raw counts ≥ 5 in at least 10% of all samples from that brain region. **g** For all brain regions and different RNA classes, we analyzed the composition of expressed RNA counts at each age point individually. The trends for all fifteen brain regions are visualized and lines are fitted with a third-degree polynomial.

**Supplementary Fig. 2: a** UMAP of all samples and for all tRNAs colored by sex. Analogue to Fig. 1d. **b** UMAP of all samples and for all tRNAs colored by age. Analogue to Fig. 1d. **c** UMAP of all samples and for all miRNAs colored by sex. Analogue to Fig. 1f. **d** UMAP of all samples and for all miRNAs colored by age. Analogue to Fig. 1f. **e** Expressed miRNAs per brain region and the overlap between them. **f** UMAP of young samples (3-, 12-, and 15-month-old) colored by brain region. **g** UMAP of all young samples according to Supplementary Fig. 2f colored by sex. **h** UMAP of the young samples according to Supplementary Fig. 2f colored by age. **i** Analogue to Fig. 2a for male and female samples combined: Heatmaps of the 50 top miRNA from all brain regions determined by coefficient of variation calculated using the medians of the expression values of each brain region. Shown are the absolute standardized expression values (z-scores). The black borders are indicating the binarization (|z-score| > 0.5) on which a clustering into four clusters using a hierarchical clustering was performed. For visualization purposes, we removed features entirely below the selected threshold. **j** Overview plot of the median expression values of mmu-miR-9-5p per brain region. Bar height represents the median value and error bar the standard deviation.

**Supplementary Fig. 3: a** Analog to Fig. 3a. Boxplots showing the other eleven trajectories of miRNA mmu-miR-9-5p. Asterisks highlight the significance of a deregulated comparison between that age and the control age (3 months). The Spearman’s rank correlation coefficient from the miRNA with age is displayed above each plot (significantly, if adjusted p-value < 0.05). **b** Spearman’s rank correlation coefficient for every miRNA with age per brain region. On the left for male samples and on the right for female samples. Colored areas highlight correlation values above 0.5 (yellow) and underneath −0.5 (green). Asterisks indicate that the corresponding adjusted p-value is significantly (smaller than 0.05). For clarity of the plot, features were not shown if there was no positive or negative correlation within a brain region. **c** Heatmaps showing for each brain region the number of significantly deregulated miRNAs for the comparison between each older age stage to 3 months (fold change ≥ 1.5 or ≤ 1/1.5, adjusted p-value < 0.05).

**Supplementary Fig. 4: a** Bar plots showing the number of significantly up- or downregulated miRNAs for at least one age comparison per brain region (fold change ≥ 1.5 or ≤ 1/1.5, adjusted p-value < 0.05). The upper bars contain miRNAs which are in the same direction significantly deregulated in more than one brain region (in yellow). The respective lower bars contain miRNAs that are unique for one brain region (colored in the corresponding brain region color). **b** Upset plot providing an overview of miRNAs changing with age per brain region and presenting the uniqueness or overlap of these candidates for each sex. To determine these candidates, we determine the Spearman’s rank correlation coefficient for each miRNA in each brain region with age and filter for significantly positively or anti-correlated features (|R| ≥ 0.5, adjusted p-value < 0.05). Additionally, we perform a DE analysis for all comparisons between each age stage to 3 months. Candidates are those features for a brain region if the feature is significantly deregulated (abs. log2-fold change ≥ log2(1.5), adjusted p-value < 0.05) for at least one comparison or significantly correlated. **c** Analogue to Supplementary Fig. 3b the Spearman’s rank correlation coefficients for all samples. Colors indicate the positive or anti-correlation and the asterisk the significance. For clarity of the plot, features were not shown if there was no positive or anti-correlation with a brain region. **d** Analogue to Fig. 3b but for all samples combined. The number of significantly positively or anti-correlated miRNAs with age per brain region (using Spearman’s rank correlation coefficient with |R| ≥ 0.5, adjusted p-value < 0.05). **e** Heatmaps showing the number of significantly deregulated miRNAs for each brain region regarding the comparison between each age stage to 3 months (fold change ≥ 1.5 or ≤ 1/1.5, adjusted p-value < 0.05). **f** Analogue to Supplementary Fig. 4a but for all samples combined. The number of miRNAs for which at least one age comparison per brain region is significantly up- or downregulated (fold change ≥ 1.5 or ≤ 1/1.5, adjusted p-value < 0.05). **g** Scatter plot of the log2-transformed fold change (comparison for every older age against 3m) against the log10-transformed median expression values without the 3 months old samples. Horizontal lines indicate the fold change thresholds at 1.5 and 1/1.5 and the vertical line is drawn at 15 rpmm. **h** Overlap between the significantly age-correlated miRNAs (positive in yellow and negative in green) from Supplementary Fig. 4c with the predicted miRNAs targeting cholinergic genes^32^. **i** Summary over all brain regions of the neighborhood of the significantly age-correlated miRNAs. For each significantly age-correlated miRNA, we accumulated the number of significantly age-correlated miRNAs within 10kb range on the same and the opposite strand. Overall brain regions we summarized these occurrences in the heatmap. If there exists no significantly age-correlated miRNA with significantly age-correlated neighbors (on the same or the opposite strand), the heatmap value is 0 and omitted for clarity.

**Supplementary Fig. 5: a** Spearman’s rank correlation coefficient values for each miRNA with age per brain region over the cumulative genomic coordinates. Yellow (green) dots indicate significantly positively (negatively) correlated miRNAs.

**Supplementary Fig. 6: a** The gene set enrichment analysis (GSEA) result created with MIEAA^54^ for six brain regions showing the top 50 (sorted by the adjusted p-value) depleted (green) and enriched (yellow) pathways over all brain regions. The analysis was performed per brain region according to the Spearman’s rank correlation coefficient direction and the obtained p-values.

**Supplementary Fig. 7: a** All cluster trajectories with a membership of at least 15% for the c-means cluster result applied to the trajectorys consisting of miRNA per brain region. All expression values are standardized and the number of clusters for a membership threshold of 15% are displayed under the cluster number.

**Supplementary Fig. 8: a** The center lines of seven clusters containing trajectories corresponding to mmu-miR-146a-5p. If trajectories for multiple brain regions were assigned to the same cluster and therefore have the same center line, we vary the line width for visualization purposes. **b** The trajectories of miRNA mmu-miR-146a-5p for all fifteen brain regions are provided. Asterisks highlight the significance of a deregulated comparison between that age and the control age (3 months). The Spearman’s rank correlation coefficient from the miRNA expression with age is displayed above each plot (significantly, if adjusted p-value < 0.05). **c** Analogue to Supplementary Fig. 8a. The center lines of eight clusters containing trajectories corresponding to mmu-miR-5100. **d** Complementing and analogue to Fig. 4h, we see the trajectories from the further eight brain regions of miRNA mmu-miR-155-5p. Asterisks highlight the significance of a deregulated comparison between that age and the control age (3 months). The Spearman’s rank correlation coefficient from the miRNA expression with age is displayed above each plot (significantly, if adjusted p-value < 0.05).

**Supplementary Fig. 9: a** Adjusted from *CNS microRNA Profiles*^39^. We display the occurrences of mmu-miR-146a-5p in different brain cell types via the fold changes between the cell type and the brainstem data. Microglia dataset: **b** Overview of the aligned (green) and not aligned (grey) reads against the mouse genome for every sample of the microglia data set. Values are given in millions. **c** Reads mapped to RNA types per sample in percentages. Ages are highlighted at the left side. Diet restriction and injection experiments: **d** Overview of the aligned (green) and not aligned (grey) reads against the mouse genome for the diet restriction experiment data set. Combined per experimental group. Values are given in millions. **e** Overview of the aligned (green) and not aligned (grey) reads against the mouse genome for the injection experiment data set. Combined per experimental group. Values are given in millions. **f** Reads mapped to RNA types per sample in percentages for the diet restriction experiment data set. Brain regions are highlighted at the left side. **g** Reads mapped to RNA types per sample in percentages for the injection experiment data set. Brain regions are highlighted at the left side. Diet restriction experiment: **h** Volcano plot presenting the results of the DE analysis for the comparison Treatment versus Control for all samples. **i** Violine plots for mmu-miR-155-5p for all fourteen brain regions visualizing the relation of the properties control and treatment (treat.). In this graphic an “Up” means that the comparison within this brain region was upregulated (fold change ≥ 1.5) and a “Down” would mean that it is downregulated (fold change ≤ 1/1.5). The horizontal line in each violin the geometric median, which was utilized to calculate the fold change. Injection experiment: **j** Volcano plot presenting the results of the DE analysis for the comparison old mice with young plasma versus PBS for all samples. Analog to Supplementary Fig. 9h. **k** Analogue to Supplementary Fig. 9i. MiR-155-5p for all eleven brain regions visualizing the relation of the properties PBS (control) and old mice with young plasma (treat.). As above an “Up” (“Down”) highlights an upregulation (downregulation) based on the fold changes. The horizontal line in each violin describes the geometric mean used for fold change calculation.

**Supplementary Fig. 10: a** Complementing and analogue to Fig. 5g for the other ten brain regions visualizing the expression of different isomiRs of mmu-miR-155. The canonical isomiR is highlighted as a dark line. **b** The scatter plots are given for five functionally validated target genes^44^ with a significant (adjusted p-value < 0.05) correlation value lower than −0.5. We see in each plot the brain regions in which this gene is significantly anti-correlated (indicated by the color) and the relation between the gene and the miR-155-5p broken down into median per brain region and age point. We used the mRNA data from Hahn et al.^3^. **c** Spearman’s rank correlation coefficient values for all genes from the mTOR pathway^45^ and mmu-miR-155-5p. Black borders indicate a correlation value below −0.3 or above 0.3 and asterisks denote a significant correlation (adjusted with Benjamini-Hochberg). The used mRNA data is from Hahn et al.^3^. **d** Intensity plot of MEF2A in hippocampus split by sex and plotted against the age. Data originates from Aging B6 Proteomics^46^. Black circles mark all data points used for the box plot.

**Supplementary Table 1:** Metadata table for brain aging, diet restriction and injection experiment and microglia datasets with all sample information containing additional alignment statistics.

**Supplementary Table 2:** Group sizes for brain aging, diet restriction and injection experiment and microglia datasets and parameters of the metadata.

**Supplementary Table 3:** Listing the miRNAs included in Fig. 2b per brain region in only male, only female or both.

**Supplementary Table 4:** Spearman’s rank correlation coefficient values and the adjusted p-values using the Benjamini-Hochberg procedure for every feature with age for male and female datasets of the aging cohort.

**Supplementary Table 5:** DE analysis results for the corresponding comparisons between the ages and per brain region. Contained are the group sizes, medians, the adjusted p-values using the Benjamini-Hochberg procedure and the fold changes for male and female datasets of the aging cohort.

**Supplementary Table 6:** A list of all age-related miRNAs obtained by male or female correlation and DE analyses.

**Supplementary Table 7:** Spearman’s rank correlation coefficient values and the adjusted p-values using the Benjamini-Hochberg procedure for every feature with age for all samples of aging cohort.

**Supplementary Table 8:** DE analysis results for the corresponding comparisons between the ages and per brain region. Contained are the group sizes, medians, the adjusted p-values using the Benjamini-Hochberg procedure and the fold for all samples of the aging cohort.

**Supplementary Table 9:** A list of all age-related miRNAs obtained by correlation and DE analyses of the complete aging cohort.

**Supplementary Table 10:** Result of the c-means clustering. Including for every trajectory composed of miRNA and brain region the corresponding cluster and the membership percentage.

**Supplementary Table 11:** DE analysis of the microglia data for the comparison old versus young.

