## Supplementary figures and images for "A spatio-temporal brain miRNA expression atlas identifies sex-independent age-related microglial driven miR-155-5p increase"

### Supplemental Figures

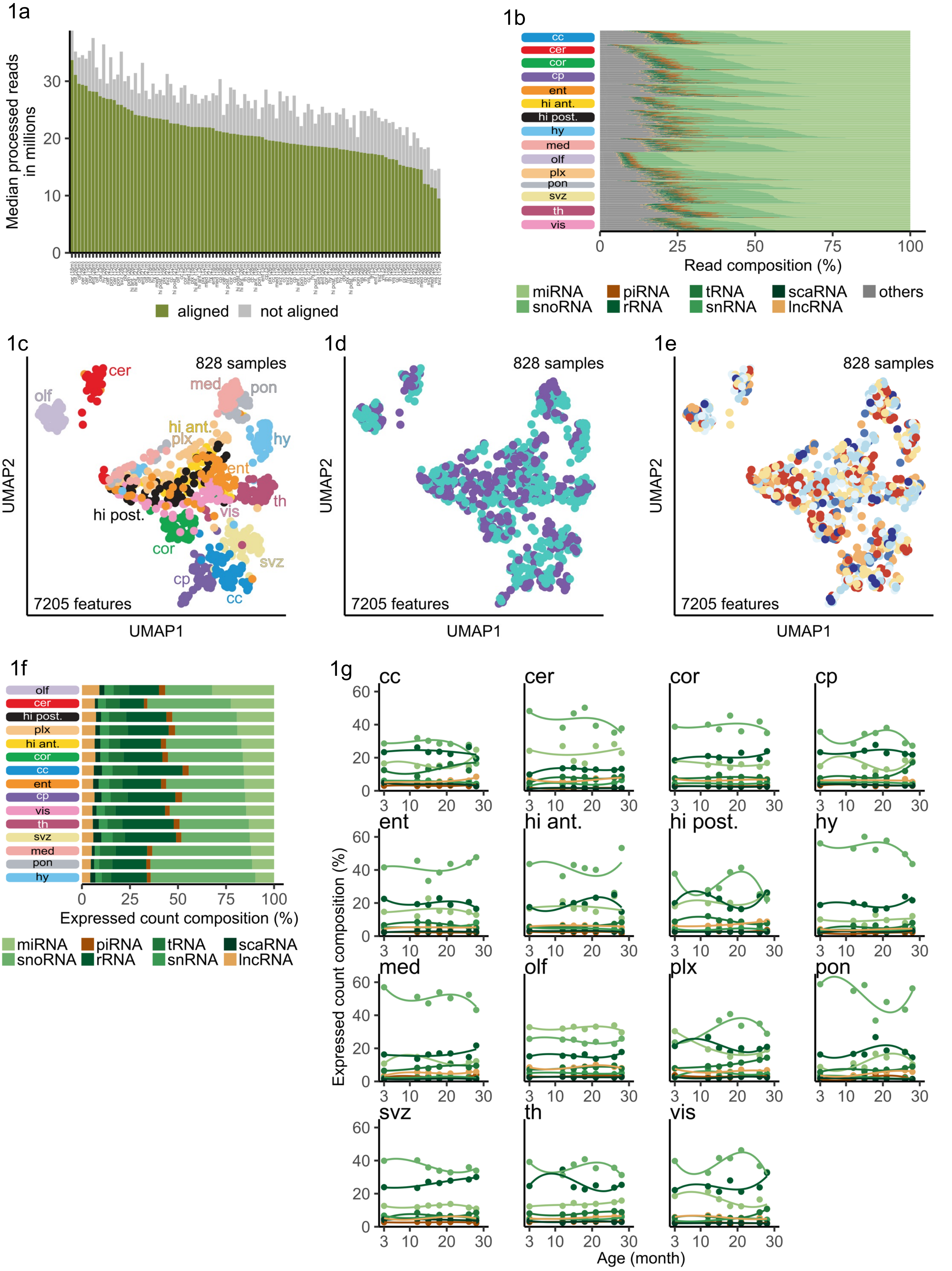

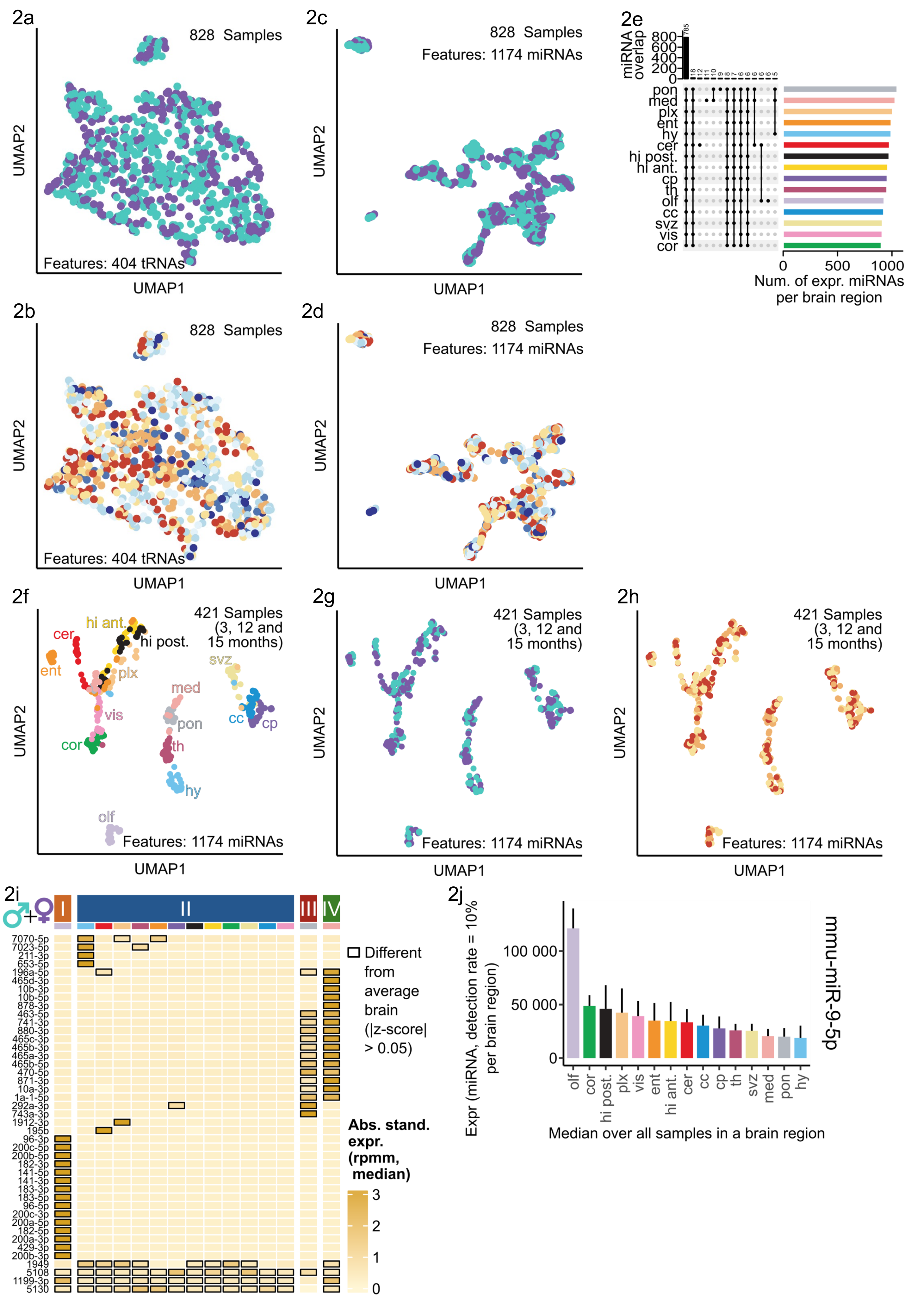

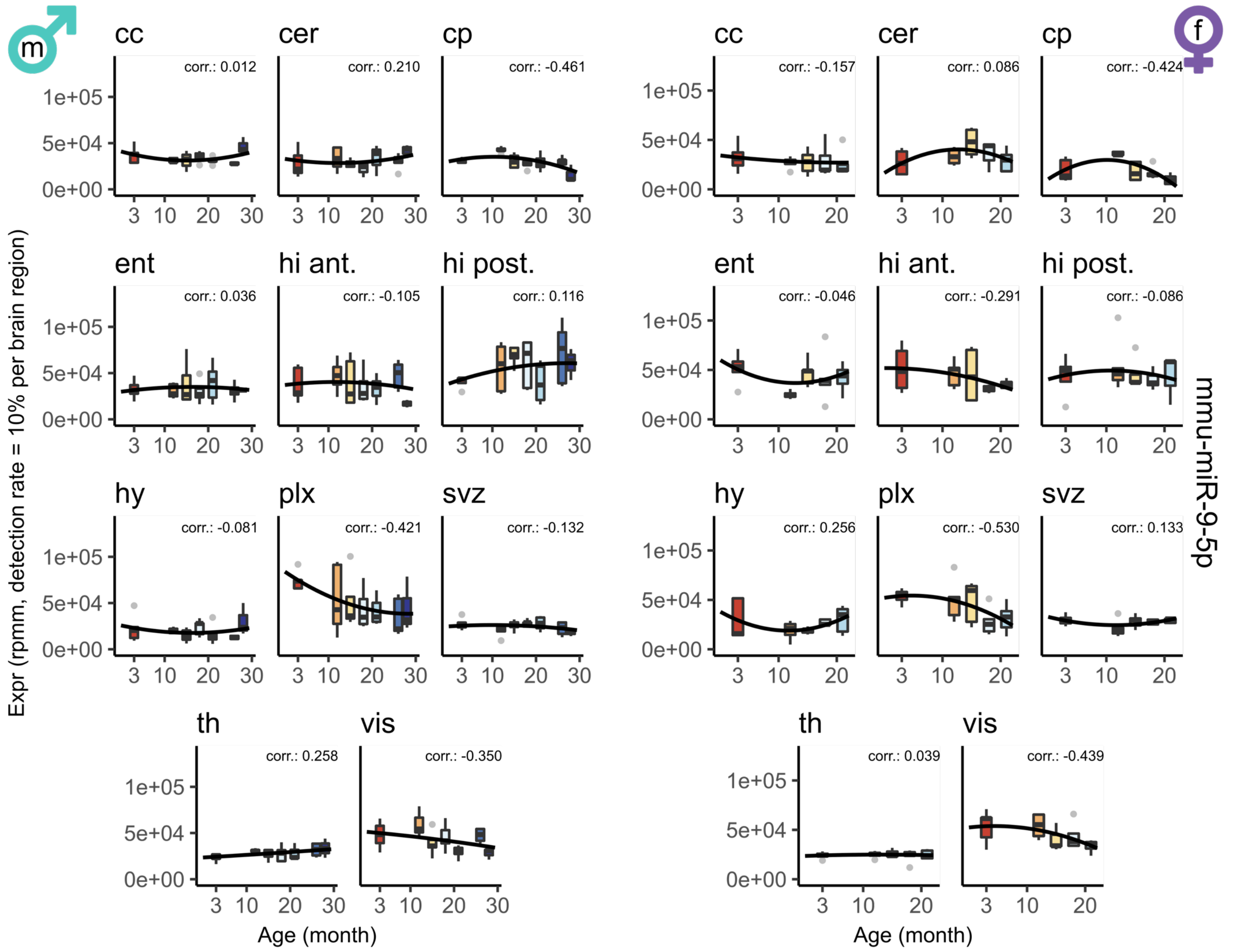

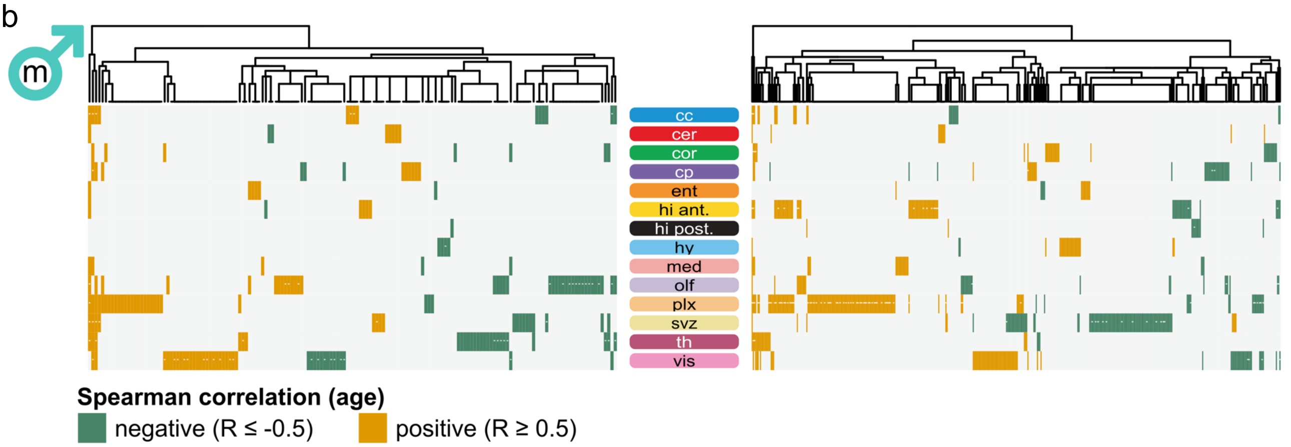

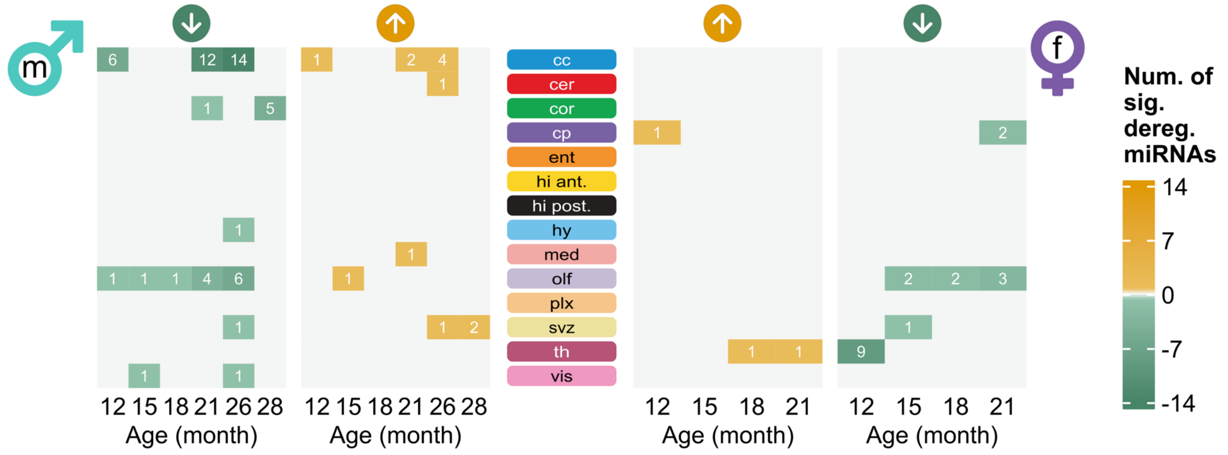

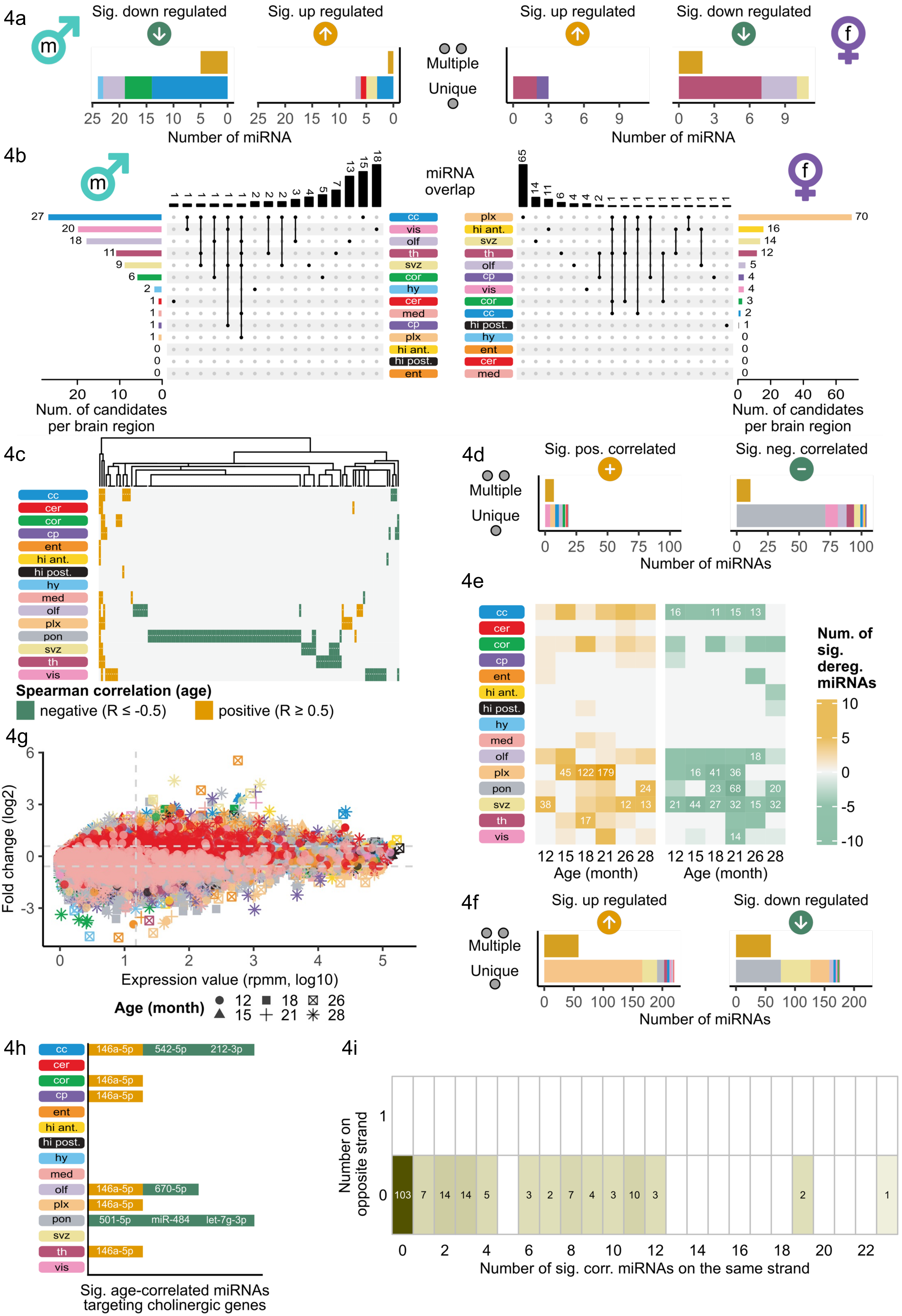

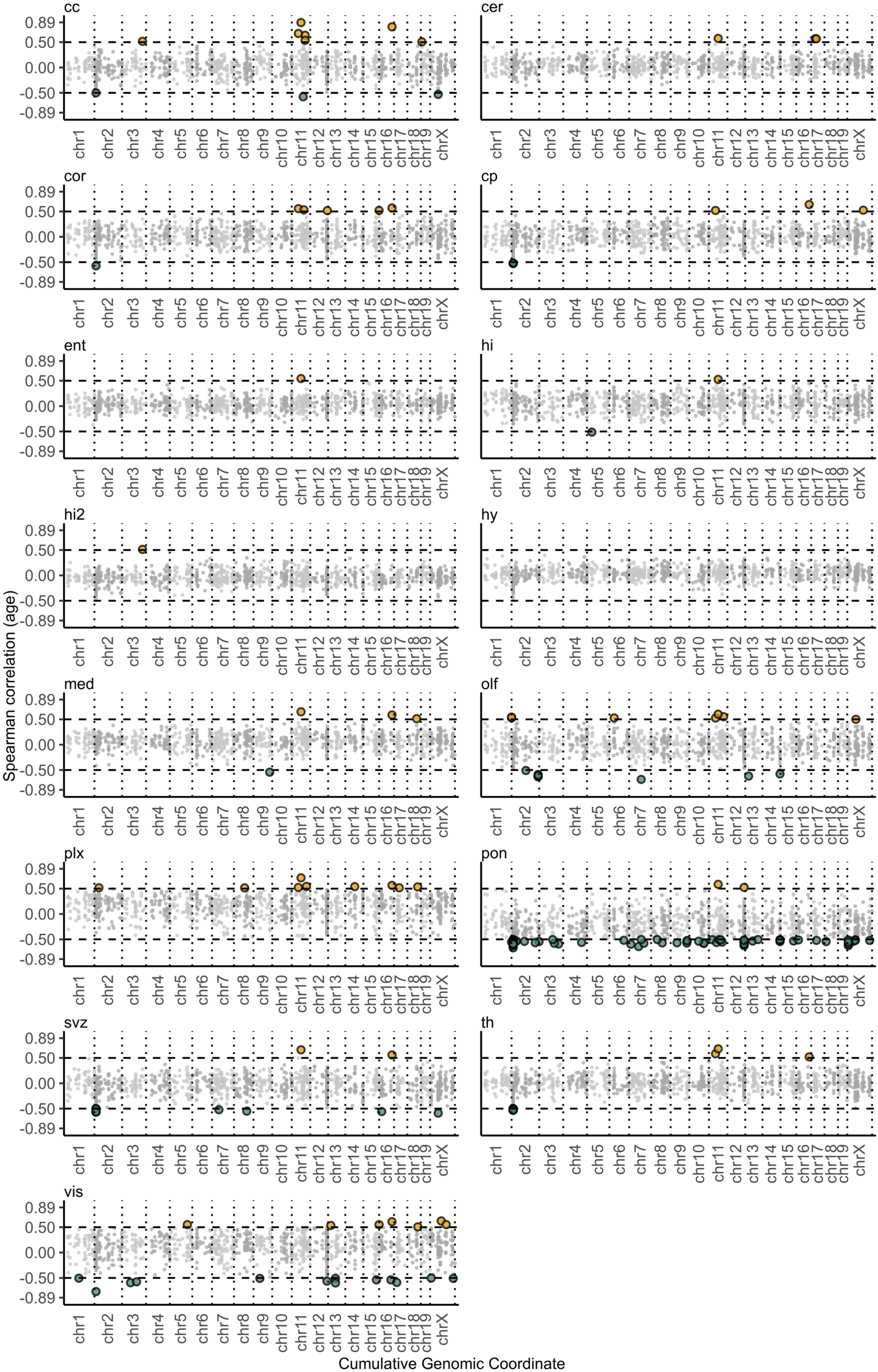

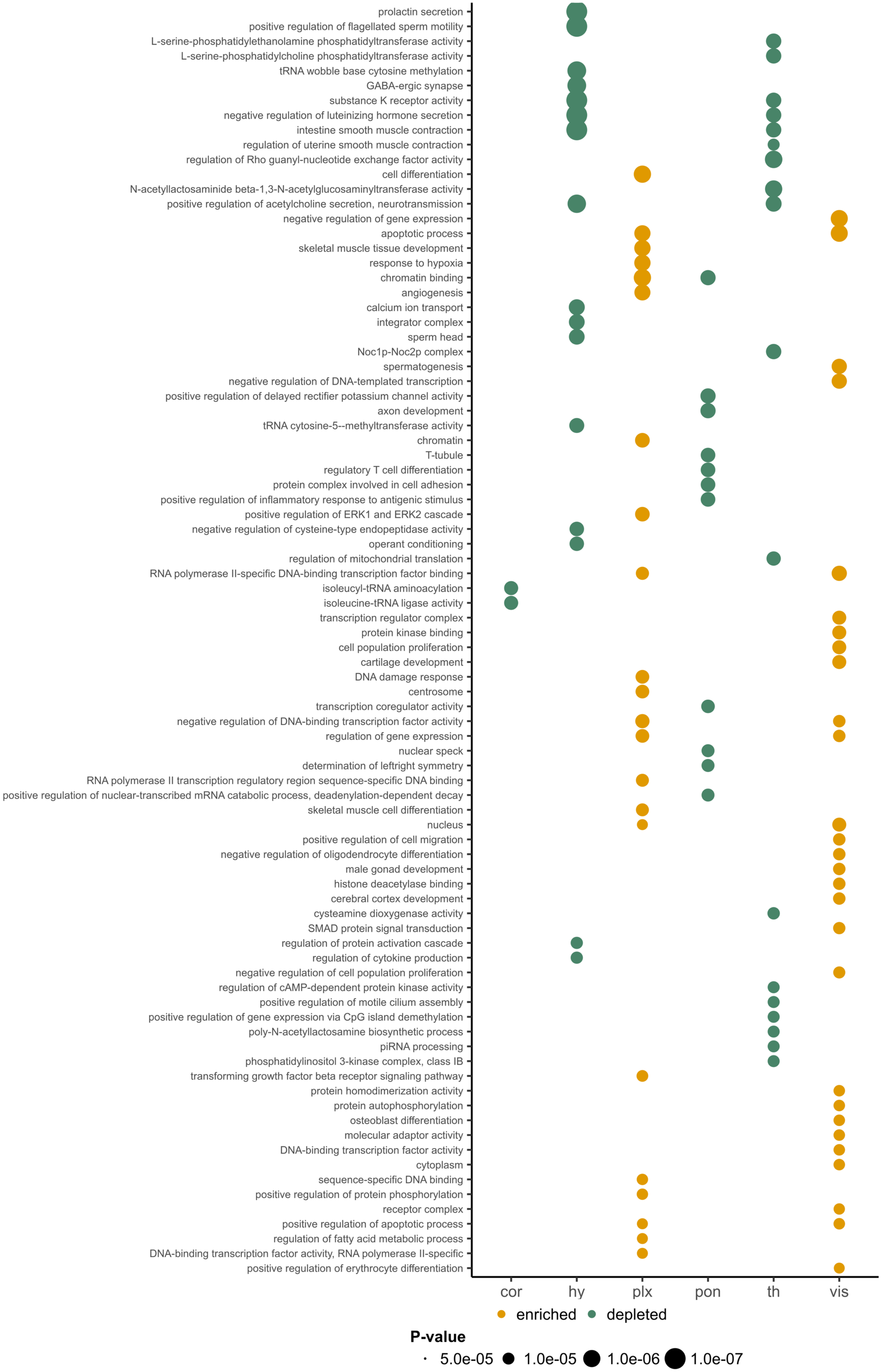

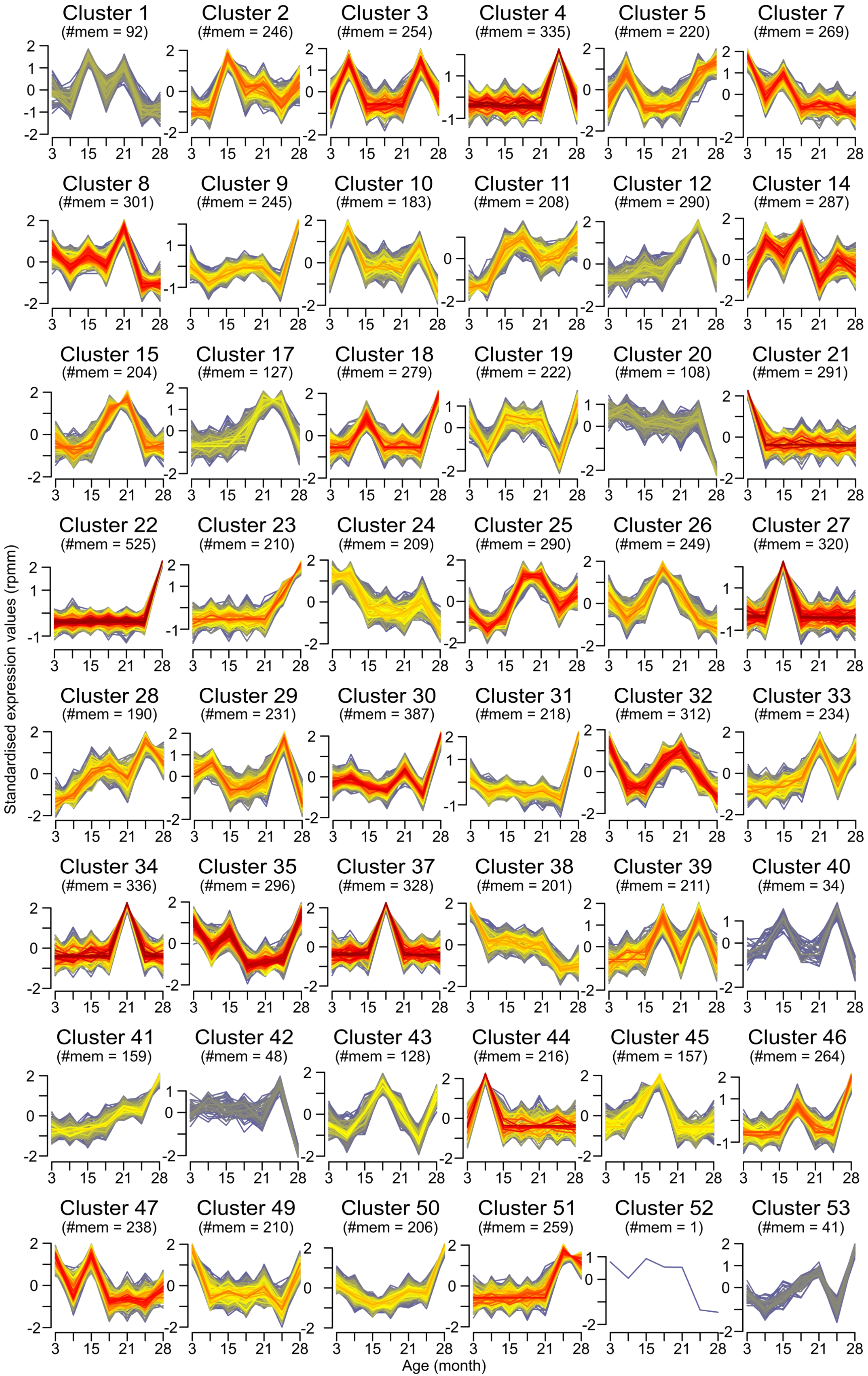

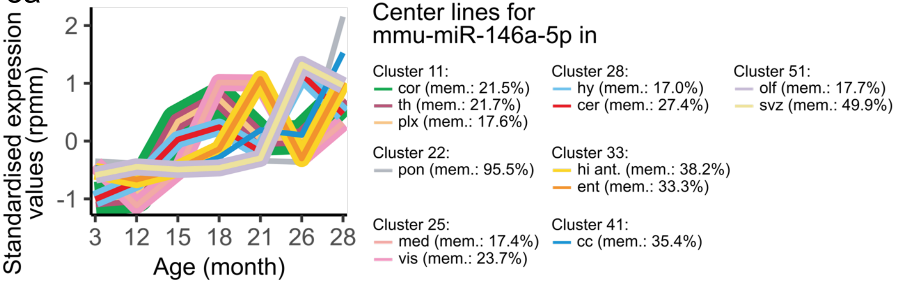

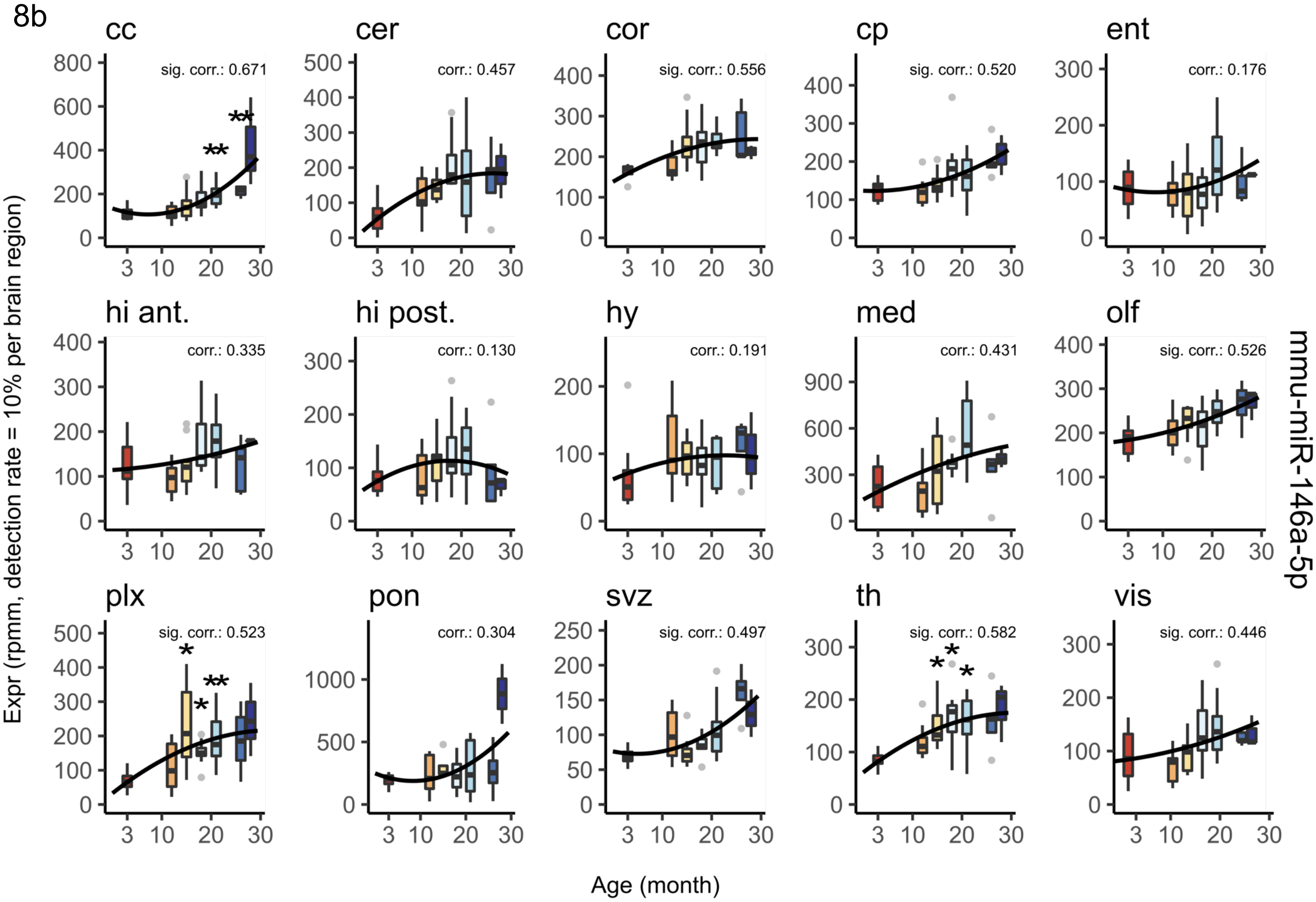

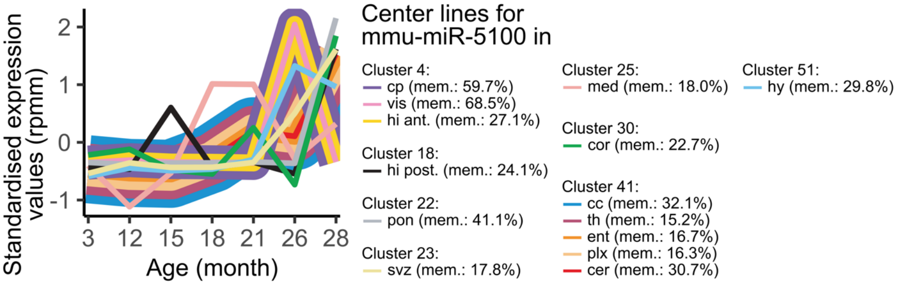

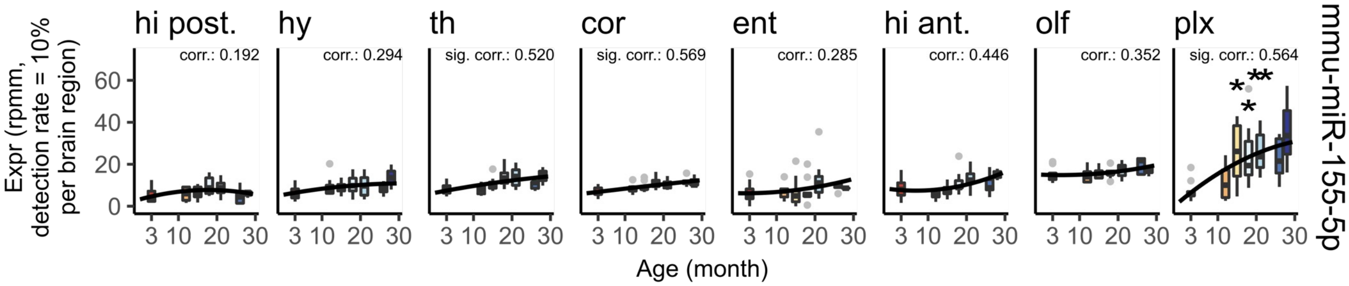

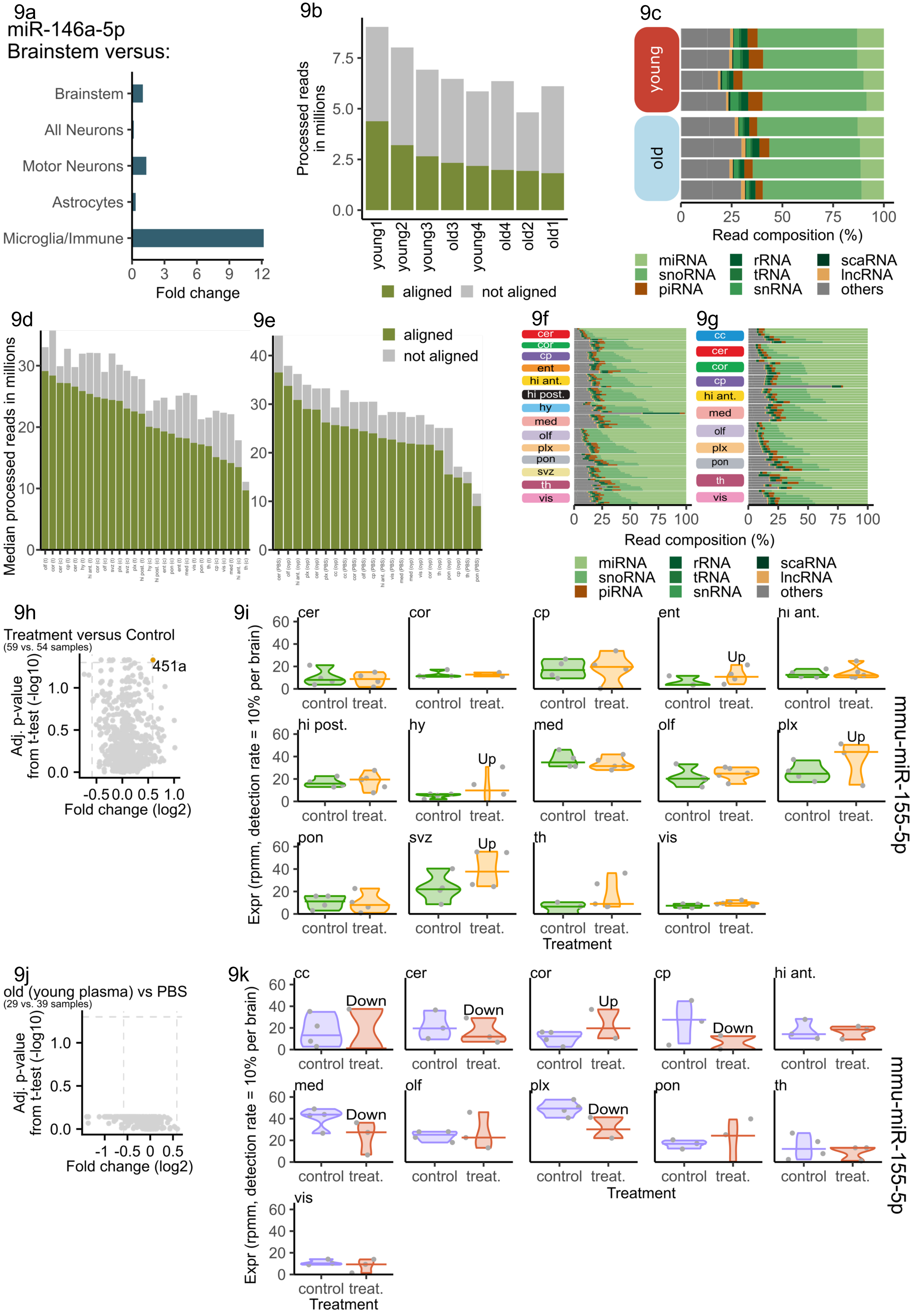

10a

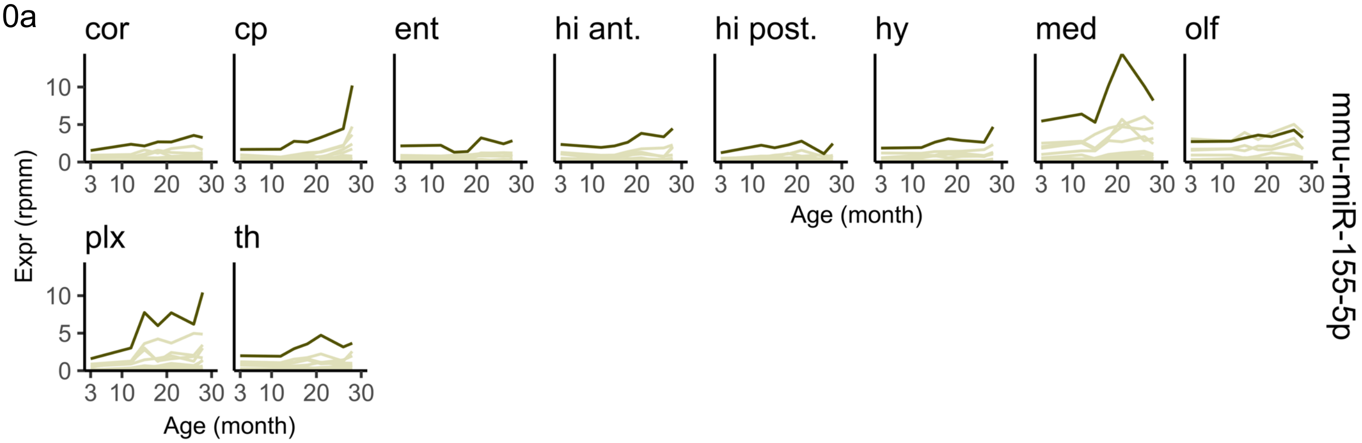


10b

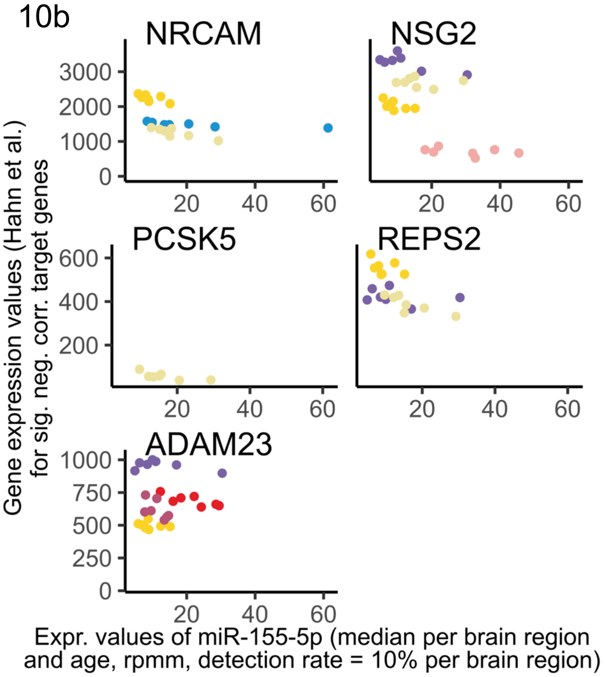


10c

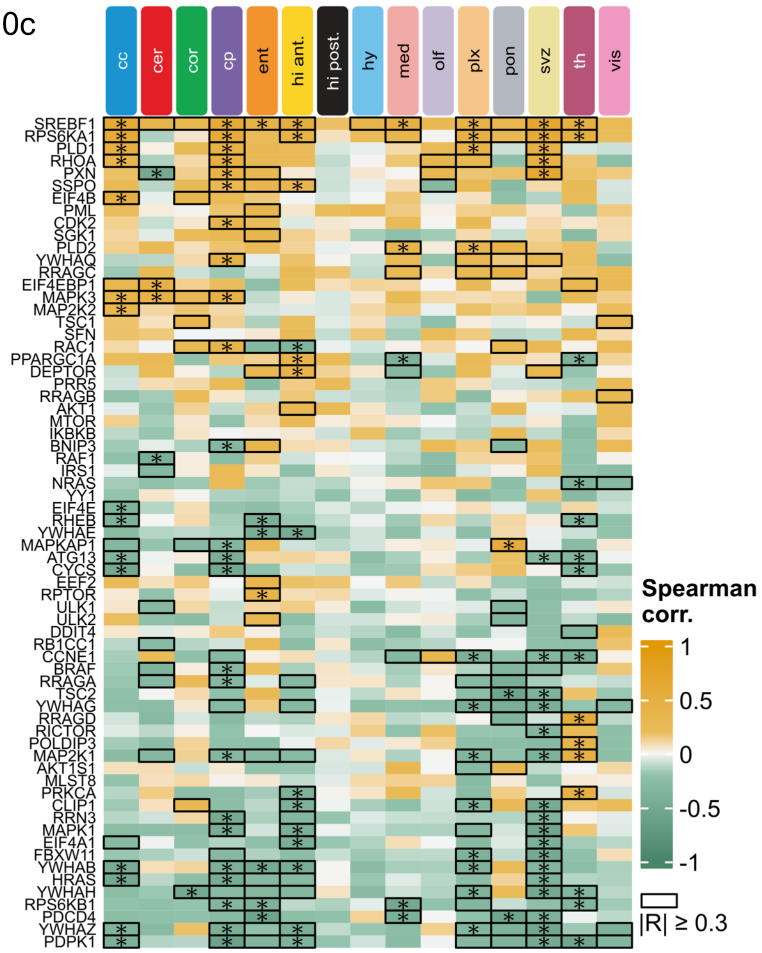


10d

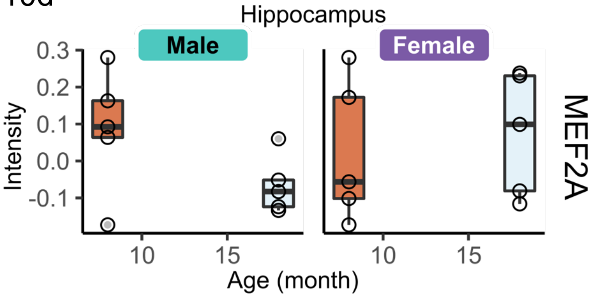
